# Quantitative single cell 5hmC sequencing reveals non-canonical gene regulation by non-CG hydroxymethylation

**DOI:** 10.1101/2021.03.23.434325

**Authors:** Emily B. Fabyanic, Peng Hu, Qi Qiu, Tong Wang, Kiara N. Berríos, Jennifer Flournoy, Daniel R. Connolly, Zhaolan Zhou, Rahul M. Kohil, Hao Wu

## Abstract

Oxidative modification of 5-methylcytosine (5mC) generates 5-hydroxymethylcytosine (5hmC), a DNA modification that exhibits unique epigenetic regulatory functions and impacts diverse biological processes. However, standard single-nucleus/cell bisulfite sequencing methods cannot resolve the base ambiguity between 5mC and 5hmC to accurately measure cell-type specific epigenomic patterns and gene regulatory functions of 5hmC or true 5mC. Here, we develop single-nucleus 5hmC sequencing (snhmC-seq) for quantitative and unbiased 5hmC profiling in single cells by harnessing differential deaminase activity of APOBEC3A towards 5mC and chemically protected 5hmC. We used snhmC-seq to profile single nuclei from cryopreserved mouse brain samples to reveal epigenetic heterogeneity of 5hmC at single-cell resolution and uncovered a non-canonical gene regulatory role of genic 5hmC in non-CG context.

## INTRODUCTION

DNA cytosine methylation is a key epigenetic modification required for proper regulation of gene expression and is dynamically regulated in a cell-type specific manner during mammalian development and organ maturation^1, 2^. Developing and adult tissues contain heterogeneous cell populations, but bulk DNA methylome profiling analysis cannot resolve epigenetic signatures contributed by distinct cell types or states^3^. Recent advances in single-cell bisulfite sequencing (BS-seq) have revolutionized the analysis of cell-type specific DNA methylomes^4–6^. However, BS-seq cannot distinguish 5mC from 5hmC^7, 8^, an oxidized form of 5mC that is generated by the Ten-eleven translocation (TET) family of DNA dioxygenases^9, 10^. As the most abundant oxidized form of 5mC, 5hmC is stably enriched at many specific genomic regions and exhibits unique epigenetic regulatory functions^11^. For instance, methyl-CpG binding protein 2 (MeCP2), is tightly bound to 5mC but not 5hmC in the CG context^12^, therefore impacting epigenome functions via its differential affinity towards 5mCG and 5hmCG. Thus, it is critical to resolve the base ambiguity between 5mC and 5hmC at single-cell levels to accurately measure cell-type specific epigenomic profiles of 5hmC or true 5mC during normal development or disease progression.

Genome-wide distribution of 5hmC in bulk samples has been profiled at single-nucleotide resolution using modified bisulfite sequencing^13, 14^, restriction enzymes^15, 16^, and recently developed bisulfite-free approaches^17–19^ (**Fig. S1**); however, most assays require ample starting material and cannot resolve epigenetic heterogeneity in 5hmC across diverse cell types or states. A recent study employed the glucosylated 5hmC (5ghmC)-dependent restriction endonuclease AbaSI to map 5hmC in single cells^20^; however, this method has several limitations. First, the restriction digestion based approach relies on selective enrichment of 5hmC-containing genomic fragments^15, 20^, therefore obscuring quantitation of absolute 5hmC levels and preventing integration with single-cell BS-seq datasets to calculate true 5mC profiles in a cell-type specific manner. Second, AbaSI exhibits preference for specific sequences and local DNA modification states^15^, presenting challenges for unbiased genomic coverage. Finally, due to the ambiguity of assigning AbaSI cutting sites within the CH context (H=A, C, and T), scAba-seq is only reliable for identifying 5hmC within CG sites^20^. Therefore, an unbiased and quantitative method is urgently needed for 5hmC profiling in single cells and for assessing regulatory functions of 5hmC across heterogeneous cell populations in primary tissues.

## RESULTS

To quantitatively profile 5hmC at base-resolution, we recently developed APOBEC-coupled epigenetic sequencing (ACE-seq)^17^, which repurposes one member of the AID/APOBEC family of DNA deaminases, APOBEC3A (A3A), for robust enzymatic deamination of unmodified C and 5mC, but not 5hmC enzymatically protected via glucosylation. In ACE-seq, DNA fragmentation, *in vitro* 5hmC glucosylation, denaturation of DNA into a single-stranded DNA (ssDNA) substrate and multiple rounds of DNA purification are required to be performed in separate steps for high-throughput sequencing (Left in **Fig. S2a**), therefore presenting challenges for genome-wide analysis of 5hmC in single cells. To streamline sample pre-processing steps, we explored bisulfite chemical conversion as a single-tube reaction to generate a preferred A3A substrate. As our prior work has suggested that A3A discriminates against cytosines with bulk 5-position substitutes ^21, 22^, we posited that bisulfite treatment could achieve chemical protection of 5hmC through formation of cytosine-5-methylenesulfonate (CMS)^8^ while simultaneously randomly fragmenting and denaturing the DNA (right in **Fig. S2a**). The bisulfite-treated, A3A-deaminated ssDNA substrates could then be efficiently captured by random priming^6^ and/or post-deamination adaptor tagging^4^. An added benefit is that bisulfite conversion would also deaminate two rare oxidized forms of 5mC, 5-formylcytosine (5fC) and 5-carboxycytosine (5caC)^23^, thus further increasing the accuracy of 5hmC profiling (right in **Fig. S2a**).

To validate this chemical/enzymatic hybrid variant of ACE-seq (termed bisulfite ACE-seq) in whole-genome, base-resolution analysis of 5hmC, we first characterized genomic DNA (gDNA) of a mutant T4 phage (∼169 kb; all cytosine bases are in 5hmC state) and *in vitro* methylated lambda phage (∼48.5 kb; enzymatically methylated at all CG sites but unmethylated at all CH sites) phage^17^. The pooled dual phage gDNA spike-ins contain known and homogenous modifications in all sequence contexts, thus providing a well-established model system for benchmarking unbiased high-throughput sequencing analysis. Whole-genome profiles of mutant T4 phage DNA showed that chemical conversion of 5hmC to CMS in bisulfite ACE-seq (5hmC in CG/CHG/CHH contexts: 98.8%/98.7%/98.8%, respectively) exhibited comparable level of protection from A3A deamination as compared to enzymatic 5hmC glucosylation in standard ACE-seq (CG/CHG/CHH contexts: 98.4%/98.5%/98.6%, respectively) (**Fig. S2b**). In analyzing methylated lambda phage gDNA, we observed robust 5mC deamination for both standard (non-conversion rate: 0.3%) and bisulfite (non-conversion rate: 0.4%) ACE-seq protocols (**Fig. S2b**), whereas the BS-seq control showed minimal conversion as expected (non-conversion rate: 98.0%). To rigorously measure the false positive rate of genome-wide 5hmC detection in standard and bisulfite ACE-seq conditions, we analyzed hypermethylated gDNA from Tet1-3 triple knockout (Tet TKO) mouse embryonic stem cells (mESCs)^24^. We observed a very low false positive rate (0.2%) in the bisulfite ACE-seq workflow, comparable to that of ACE-seq (0.3%) (right in **Fig. S3a**).

While separate DNA shearing and heat denaturing steps are required by standard ACE-seq, we reasoned that simultaneous fragmentation and denaturation of gDNA during the initial bisulfite chemical reaction would render these sample processing steps optional in bisulfite ACE-seq. Thus, a small number of nuclei isolated from cryo-preserved primary tissues could be directly analyzed using bisulfite ACE-seq, expanding the utility of this approach for analyzing precious clinical samples. To test this, we analyzed whole-genome 5hmC profiles in adult neuronal nuclei that are known to be enriched for 5hmC^17^ (left in **Fig. S3a**). First, whole-genome 5hmC profiles from bisulfite and standard ACE-seq were strongly correlated (Pearson’s *r*=0.93 in **Fig. S3b**). Second, compared to standard ACE-seq analysis of wild-type (WT) neuronal and Tet TKO mESC gDNA (**Fig. S3b**), bisulfite ACE-seq generated highly concordant and specific whole-genome 5hmC profiles across a wide range of input amounts (100 versus 10,000 nuclei), and inclusion of a separate heat denaturing step was dispensable, post bisulfite conversion (**Fig. S3a** and **S3c**). Third, analysis of internal spike-in phage gDNA (methylated lambda and mutant T4 phages) further confirmed robust 5hmC discrimination in these experimental conditions (**Fig. S3d**). Finally, we performed integrated (hydroxy)methylome analysis of excitatory neuronal gDNA using BS-seq (mapping 5mC+5hmC) and bisulfite ACE-seq protocols (**Fig. S4a-b**). Paired analysis of the same pool of bisulfited converted gDNA enabled highly reproducible profiling of 5mC and 5hmC across neuronal genomes (**Fig. S4c**) and uncovered single-base resolution patterns of true 5mC after subtracting bisulfite ACE-seq signals from BS-seq signals (**Fig. S4d**). Collectively, these data demonstrate that bisulfite ACE-seq, which integrates a single-tube bisulfite chemical reaction with A3A enzymatic deamination of 5mC, can accurately profile 5hmC at base-resolution across the complex mammalian genome in low input gDNA or a small number of nuclei isolated from cryo-preserved primary tissues.

To harness the robust performance of bisulfite ACE-seq to achieve quantitative and unbiased 5hmC profiling in single cells, we adopted the workflow of single-nucleus methylcytosine sequencing (snmC-seq)^4^ (**Fig. S5a**), which enables scalable generation of high quality single-cell DNA methylomes (mapping 5mC+5hmC) and has been deployed in several large-scale single-cell epigenomic profiling projects^4, 25^. In this single-nucleus hydroxymethylcytosine sequencing (snhmC-seq) strategy (**Fig. 1a**), single cells or nuclei are first sorted into 96-well plates for a one-pot bisulfite conversion reaction (gDNA fragmentation/denaturation and 5hmC protection via CMS formation) and, using A3A reaction conditions optimized for bisulfite converted gDNA from directly lysed nuclei (**Fig. S5b**), both C and 5mC are robustly deaminated. Next, the first level of sample indexing (via inline barcodes) is added by performing random primer extension of bisulfite- and A3A-treated single-stranded DNA (ssDNA) fragments (**Fig. 1a**). After pooling of pre-indexed samples (up to 8 in this study) and adaptor ligation, the second level of dual-index barcodes is incorporated through PCR amplification (**Fig. S5a** and **S6**). Finally, the amplified and indexed samples are pooled and subjected to high throughput pair-end sequencing. We note that the scalability of snhmC-seq is tunable by the number of inline (random priming), dual indexing (PCR amplification) barcodes and the use of a liquid handler^26^.

**Figure 1:**
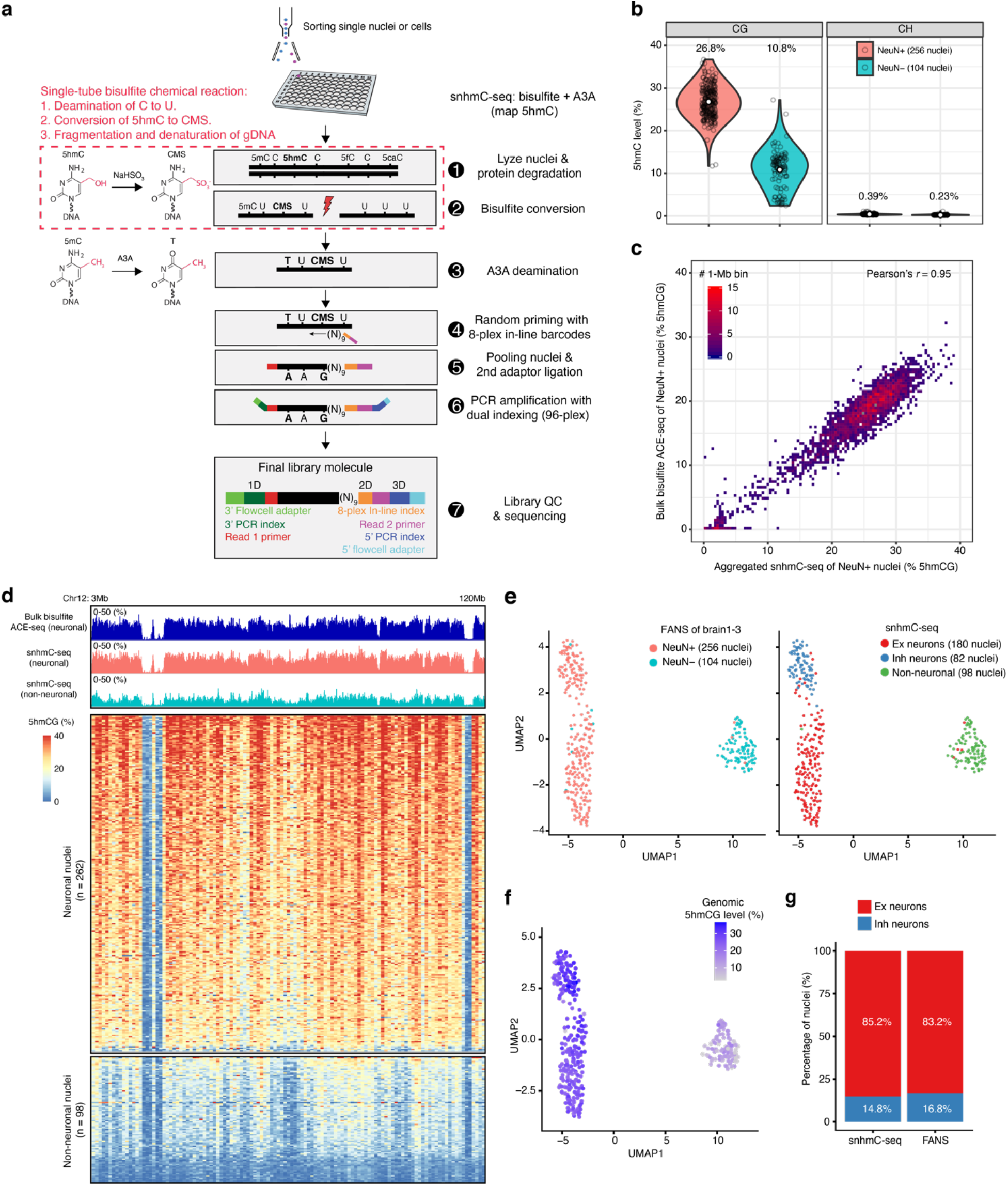
snhmC-seq workflow and performance. **a,** The snhmC-seq workflow. **b,** Violin plots comparing global CG or CH 5-hydroxymethylation levels between neuronal (NeuN+, n = 256) and non-neuronal (NeuN-, n = 104) nuclei, with mean values above each plot. The width of the violin plot corresponds to the kernel probability density of the data at a given value, and the white circle indicates mean value in each group. **c,** Correlation density plot between neuronal 5hmCG signals (averaged within 1-Mb bins spanning the genome) in bulk bisulfite ACE-seq and aggregated snhmC-seq results. **d,** Shown are the 5hmCG levels across chromosome 12 from bulk ACE- seq and snhmC-seq data sets. Upper panels show genome browser tracks of the bulk neuronal 5hmC profile (from bisulfite ACE-seq) and aggregated single-cell 5hmC profiles in NeuN+ or NeuN- nuclei (from snhmC-seq). The heatmap in the bottom panel shows the 5hmCG level in 1 Mb bins (column) across 360 single nuclei (row). The 5hmCG level is scaled by colors. Maximum level was set to 40%. **e,** UMAP visualization of clustered mouse cortical nuclei colored by FANS sorting channels (left) or cell types defined by single nucleus 5hmC profiles (right). Ex: excitatory neurons; Inh: inhibitory neurons. **f,** UMAP visualization (same as **e**) colored by global 5hmCG level. **g**, Bar plots showing the percentage of excitatory (Ex) and inhibitory (Inh) neurons identified by snhmC-seq (left) or FANS (right).

To evaluate the ability of snhmC-seq to detect cell-type specific 5hmC profiles, single nuclei labeled with a NeuN antibody (NeuN+: neuronal nuclei; NeuN-: non-neuronal nuclei) were isolated by fluorescence-activated nucleus sorting (FANS) from the mouse cortex (**Fig. S7a**). The specificity of the NeuN antibody in FANS (99.7%) was independently verified using single-nucleus RNA-seq (sNucDrop-seq^27^) analysis of NeuN+ and NeuN- nuclei (**Fig. S7b**). From three biologically independent samples (brain 1-3), we generated genome-wide 5hmC profiles in 360 single cells (**Table S1**). We observed a mean alignment rate of 46 ± 9% for our snhmC-seq assay, which is comparable to that of snmC-seq (50 ± 7%)^4^ (**Fig. S8a**). Furthermore, there were no strong biases in mapping rate across random primers with distinct inline barcodes or for specific cell types (**Fig. S8b**). At a mean sequencing depth of ∼1.36 million uniquely aligned reads per cell (on average 515,779 high-quality reads retained per cell after removing low quality and PCR duplicated reads), snhmC-seq achieved a mean coverage of 474,072 CG and 11,280,623 CH sites per cell. As a result, the coverage of mappable CG sites ranges from 0.43% to 7.1% (mean 2.4 ± 1.2%). We note that increased sequencing effort is likely to augment genomic coverage, as libraries were not near saturation (**Fig. S8c**), and the performance of snhmC-seq in mapping cytosine sites is highly similar to that of snmC-seq at matching sequencing depth (**Fig. S8d**). We next summarized mean cytosine modification levels for each cell. Compared to snmC-seq analysis (CG: 74.5%, CH: 2.5% in NeuN+ nuclei), which reveals the sum of both 5mC and 5hmC levels, snhmC-seq specifically uncovered the 5hmC level in NeuN+ nuclei (**Fig. S8e**). The averaged 5hmCG and 5hmCH levels in NeuN+ cells (CG: 26.8%, CH: 0.39%) are markedly higher than those in NeuN- cells (CG: 10.8%, CH: 0.23%) (**Fig. 1b**), in line with a previously reported bulk 5hmC analysis in the human cortex ^28^. Finally, the aggregated 5hmCG profiles of single nuclei (129 NeuN+ nuclei from brain 1 and 2) correlated strongly with that of bulk assay analysis of 10,000 NeuN+ nuclei (Pearson’s *r*=0.95 in **Fig. 1c**). At the single-cell level, the neuronal 5hmCG profiles determined by snhmC-seq (heatmap in **Fig. 1d**) tracked closely with that of the bulk dataset (genome browser view in **Fig. 1d**). Thus, snhmC-seq permitted quantitative and genome-wide profiling of 5hmC in single cells.

To evaluate whether single cell 5hmC profiles can be used for cell-type discrimination, we calculated the 5hmCG level for each cell in nonoverlapping 100-kb bins across the genome, followed by dimensionality reduction and visualization using the uniform manifold approximation and project (UMAP). We observed that snhmC-seq analysis readily separated NeuN+ neurons from NeuN- non-neuronal cells with high accuracy (left in **Fig. 1e**), which is likely due to a notable (∼2.5-fold) global difference in 5hmC levels between neuronal and non-neuronal cell populations (**Fig. 1b** and **1f**). To test whether single cell 5hmC profiling can distinguish neuronal subtypes (Ex: excitatory neurons versus Inh: inhibitory neurons), we isolated Ex (NeuN+/Neurod6+) and Inh (NeuN+/Neurod6-) nuclei using FANS from a *Neurod6/NEX*-Cre driven transgenic reporter mouse line^29^ (**Fig. S7a** and brain3 in **Fig. S9a**). Unbiased clustering analysis of NeuN+ nuclei (brain1 and 2 in **Fig. S9a**) revealed two major cell clusters corresponding to Ex (85.2%) and Inh (14.8%) neurons (right in **Fig. 1e**), which are comparable to the relative composition of Ex (83.2%) and Inh (16.8%) neurons that were prospectively determined by FANS (**Fig. 1g**). Using neuronal subtype assignment by FANS as the ground truth, the accuracy of snhmC-seq based clustering was determined to be 93.7% (119 out of 127 nuclei from brain3 in **Fig. S9a**). Notably, the single cell 5hmC profile-based clustering was not strongly associated with a number of experimental (**Fig. 1f** and left in **Fig. S9a**) and analysis parameters (**Fig. S9b**), indicating the robustness of the clustering analysis.

Recent single-cell DNA methylome (snmC-seq) analysis of the mouse cortex demonstrated that gene body CH methylation (average 5mCH+5hmCH across the annotated genic region) is inversely correlated with gene expression^4^, and identity of neuronal subtypes can be accurately annotated on the basis of depletion of genic CH methylation at known cortical neuronal subtype markers^4^. This relationship could also be leveraged for computationally integrating single-cell DNA methylome data with single-cell/nucleus RNA sequencing data^30^. To integrate single-cell 5hmC profiling datasets from snhmC-seq with the single-cell DNA methylome, we first explored the relationship between 5hmC (bisulfite ACE-seq) and 5mC+5hmC (BS-seq) across the genome in the paired bulk assays (**Fig. S4a-b**). Interestingly, we observed a strong positive correlation between 5hmCH and 5mCH+5hmCH (Pearson’s *r*=0.90, middle in **Fig. S4e**), whereas 5hmCG was weakly anti-correlated with 5mCG+5hmCG (Pearson’s *r*=-0.16, left in **Fig. S4e**). In concordance with previous reports using orthogonal bulk 5hmC profiling assays ^17, 28^, 5hmCH levels (range: ∼0.2-0.6%) are substantially lower than 5mCH levels (range: ∼0.3-3.5%) in neurons (**Fig. S4e**), but neuronal 5hmCH signals are highly specific when compared to a constant baseline signal of ∼0.2% detected in Tet TKO mESCs (right in **Fig. S4e**).

We next jointly analyzed gene body 5hmCH (snhmC-seq) and CH methylation (5mCH+5hmCH in snmC-seq)^4^ for computational integration of the two datasets (see **Methods**). Using prospectively isolated Ex and Inh neurons (from brain 3) as ground truth controls, we observed that 94.7% Ex (54 out of 57 Ex nuclei defined by FANS) and 84.3% Inh (59 out of 70 Inh nuclei defined by FANS) neurons were correctly assigned to neuronal subtypes previously annotated by snmC-seq analysis of the adult mouse cortex. The overall concordance of Ex and Inh neuronal subtype assignment between snhmC-seq and snmC-seq datasets is ∼85% (152/180 Ex nuclei in snhmC-seq, 70/82 Inh nuclei in snhmC-seq) (**Fig. 2a**). The global levels of 5hmC (snhmC-seq) and 5mC+5hmC (snmC-seq) were partially concordant across neuronal subtypes in CG or CH contexts (**Fig. S10a**). Thus, these results indicate that genic 5hmCH signals (similar to CH methylation) are informative for inferring cell-type specific gene expression levels and cellular identity.

**Figure 2:**
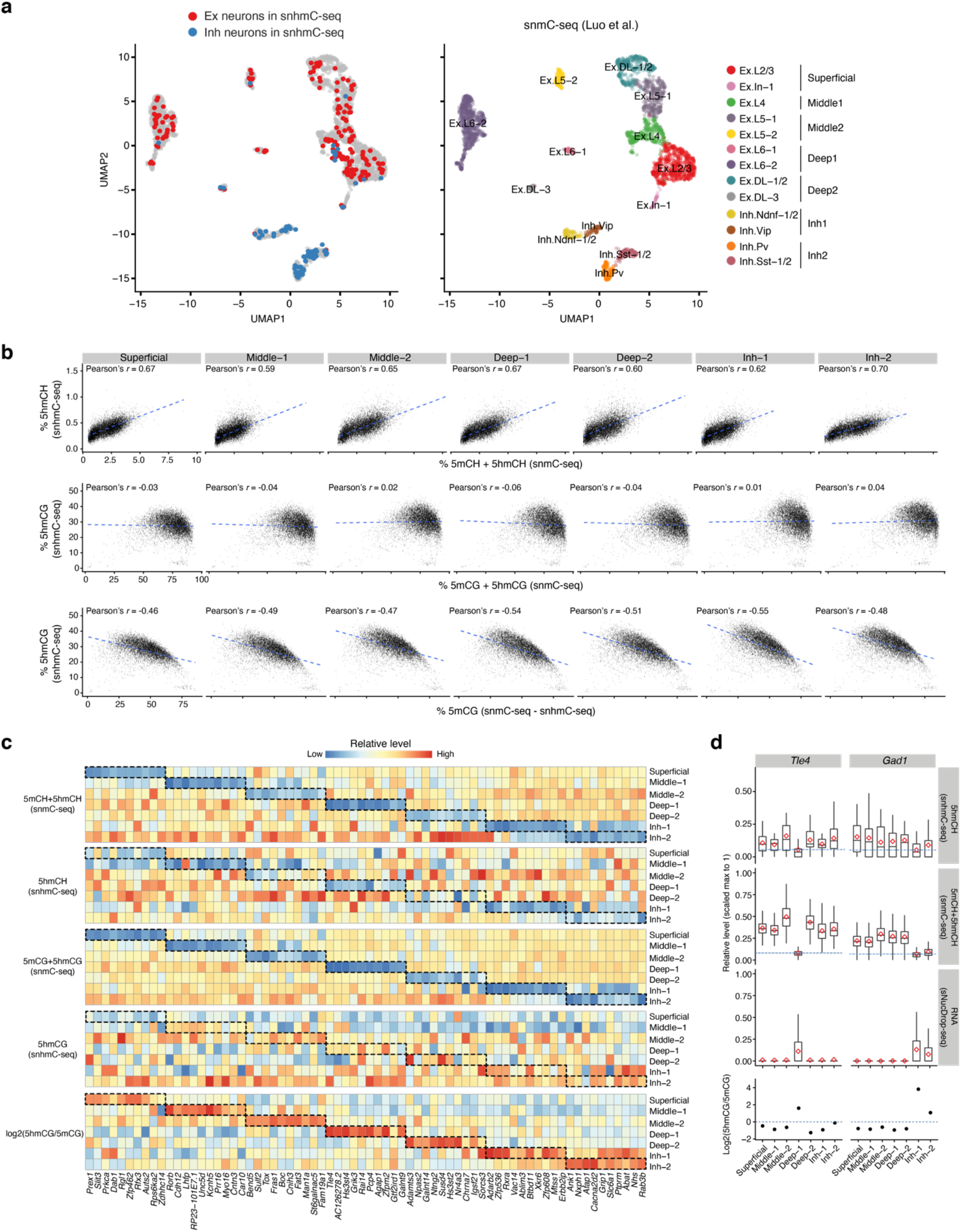
Joint analysis of snhmC-seq and snmC-seq reveals 5hmC and true 5mC profiles across diverse neuronal subtypes. **a,** UMAP visualization of mouse cortical neuronal subtypes in the adult cortex, colored by cell-type annotations defined by snhmC-seq (left panel, n = 262 Ex/Inh nuclei) or by snmC-seq (right panel, n = 3,377 NeuN+ nuclei). Ex: excitatory, Inh: inhibitory, DL: deep layer. Cell-type annotations were simplified by combining subtypes within the same anatomical cortical layer (Superficial, Middle1/2 and Deep1/2) or according to their transcriptomic similarity (for Inh1/2). **b,** Scatterplots comparing gene-body 5hmCH from snhmC-seq with genic CH methylation (5mCH+5hmCH) from snmC-seq (upper panel), 5hmCG with CG methylation (5mCG+5hmCG) (middle panel), and 5hmCG with true 5mCG (bottom panel) in 7 major neuronal subtypes. True 5mCG was calculated by directly subtracting 5hmCG level measured by snhmC- seq from 5mCG+5hmCG level measured by snmC-seq. The linear regression is indicated by a blue dashed line and Pearson’s correlation coefficient is shown above the plot. **c,** Heat maps of genic CH methylation (5mCH+5hmCH), 5hmCH, CG methylation (5mCG+5hmCG), 5hmCG, log2-scaled ratio of 5hmCG and true 5mCG levels for the top 10 cell-type specific markers defined by depletion of genic CH methylation level across 7 neuronal subtypes. The average methylation level was calculated in each defined neuronal subtype, then was centered and scaled gene-wise. **d,** Joint analysis of 5hmCH from snhmC-seq, 5hmCH + 5mCH from snmC-seq and mRNA expression from sNucDrop-seq in neuronal subtypes. Box plots (upper panel) showing the scaled level of 5hmCH, 5hmCH and mRNA expression (transcripts per 10K) of two representative neuronal subtype specific genes, *Tle4* and *Gad1*. The red diamond in the box indicates mean value of each group. The lowest average level of measurements in the box plots is indicated by blue dashed lines. Also shown in the bottom panel are the log2-scaled ratios between 5hmCG and true 5mCG levels. The ratio of 1 is indicated by the blue dashed line. See ‘Data visualization’ in the Methods for definitions of box plot elements.

Previous bulk DNA methylome profiling studies in FANS-isolated specific neuronal populations indicate that intragenic 5mCG and 5hmCG levels are negatively and positively correlated with gene expression in the same cell population, respectively ^17, 28^; however, the cell-type specific gene regulatory role(s) of 5hmC in non-CG context remains largely unclear. To address this, we first analyzed the relationship between gene body 5mC and 5hmC in both CG and CH contexts across different neuronal sub-types. Cell-type specific 5hmC profiles revealed that genic 5hmCH is positively correlated with CH methylation (5mCH+5hmCH) in all cortical neuronal subtypes (top in **Fig. 2b**), whereas genic 5hmCG is negatively correlated with true 5mCG profiles (bottom in **Fig. 2b**), but not with 5mCG+5hmCG profiles (middle in **Fig. 2b**), within the same neuronal subtype. Thus, joint epigenomic profiling analyses of (hydroxy)methylomes not only confirm that genic 5hmCG (generally associated with active transcription) is functionally antagonistic to 5mCG mediated repression, but also raises the possibility that genic 5hmCH, similar to 5mCH, is associated with transcriptional repression of neuronal sub-type specific genes.

To further analyze the relationship between genic 5hmCH and gene expression, we computationally integrated single-nucleus epigenomic profiles (snhmC-seq/snmC-seq) with transcriptomic profiles (sNucDrop-seq) (**Fig. S10b**). This analysis revealed that, among top cell-type specific markers defined by low genic CH methylation in snmC-seq, gene body 5hmCH was generally depleted, whereas functional DNA demethylation via oxidizing 5mCG to 5hmCG (measured by 5hmCG/5mCG ratios)^31^, but not absolute level of genic 5hmCG, was positively correlated with transcriptionally active state (**Fig. 2c**). Indeed, down-regulation of genic 5hmCH is specifically associated with transcription of cell-type specific markers (**Fig. 2d**). Taken together, our single-cell 5hmC sequencing analysis provides strong *in vivo* evidence supporting that in contrast to the known role of 5hmCG in functionally opposing repressive 5mCG, genic 5hmCH may negatively impact the transcription of cell-type specific genes in a wide-range of neuronal sub-types.

## DISCUSSION

Given that asymmetric CH sites are much more abundant (∼1.1 billion in mouse genome) than CG sites (∼22 million symmetric CpG dyads)^32^, even a small fraction of modified 5hmCH sites (0.2-0.6%) may lead to a considerable impact on epigenetic gene regulation. Although the exact mechanism downstream of 5hmCH is currently unclear, this non-canonical gene regulatory function of 5hmCH is reminiscent of the repressive role of 5mCH ^33^ and may possibly be mediated by recruiting transcriptional repressor MeCP2, as previously predicted by *in vitro* biochemical binding assays ^12, 34^. Intriguingly, a recent study of maturing cerebellar Purkinje neurons also shows that accumulation of both 5mCH and 5hmCH (measured by bulk BS-seq and oxBS-seq assays) within gene body of a cohort of developmentally down-regulated genes is accompanied by elevated level of repressive histone modification H3K27me3 (deposited by Polycomb repression complexes)^35^, suggesting an alternative but non-mutually exclusive transcriptional inhibition mechanism mediated by genic 5mCH and 5hmCH.

In summary, our work demonstrates that snhmC-seq is a powerful approach to quantitatively measure 5hmC across genomes of single cells from heterogenous primary tissues and to reveal distinct gene regulatory functions of 5hmC in specific sequence contexts. Given its robust performance and relative simplicity, snhmC-seq has the potential to become the method of choice for high-throughput 5hmC sequencing analysis in rare cell populations or in single cells. Notably, other single-cell BS-seq protocols that leverage either a post-bisulfite adaptor tagging strategy (scBS-seq)^6^ or C-depleted adaptors (sci- MET)^5^ can also be readily modified to leverage bisulfite ACE-seq chemistry for profiling 5hmC in single cells. We envision that integrated single-cell analyses of DNA (hydroxy)methylomes can systematically determine the cell-type specific genomic patterns of 5hmC and true 5mC and reveal gene regulatory roles of these epigenetic modifications across diverse cell types or states present in normal development and human diseases.

## ACKNOWLEDGEMENTS

We are grateful to all members of the Wu and Kohli lab for helpful discussion. This work was supported by the Penn Epigenetics Institute pilot grant, the National Human Genome Research Institute (NHGRI) grant R01-HG010646 (to R.M.K. and H.W.). H.W. is also supported by National Heart Lung and Blood Institute (NHLBI) grant DP2-HL142044 and National Cancer Institute (NCI) grant U2C-CA233285.

## AUTHOR CONTRIBUTIONS

E.B.F. and H.W. conceived of and developed snhmC-seq. E.B.F. conducted most of the experiments. K.N.B., Q.Q. and J.F. performed A3A protein purification and characterization. Q.Q. and E.B.F. performed sNucDrop-seq experiments. T.W. and R.M.K. contributed reagents and helped with the experimental design. E.B.F., D.R.C., and Z.Z. performed neuronal nuclei isolation. P.H. and H.W. performed the computational analysis. P.H. developed the computational pipeline for snhmC-seq analysis with input from H.W. E.B.F., P.H., and H.W. analyzed the results and wrote the manuscript, with contributions from all of the authors.

---

**Figure S1:**
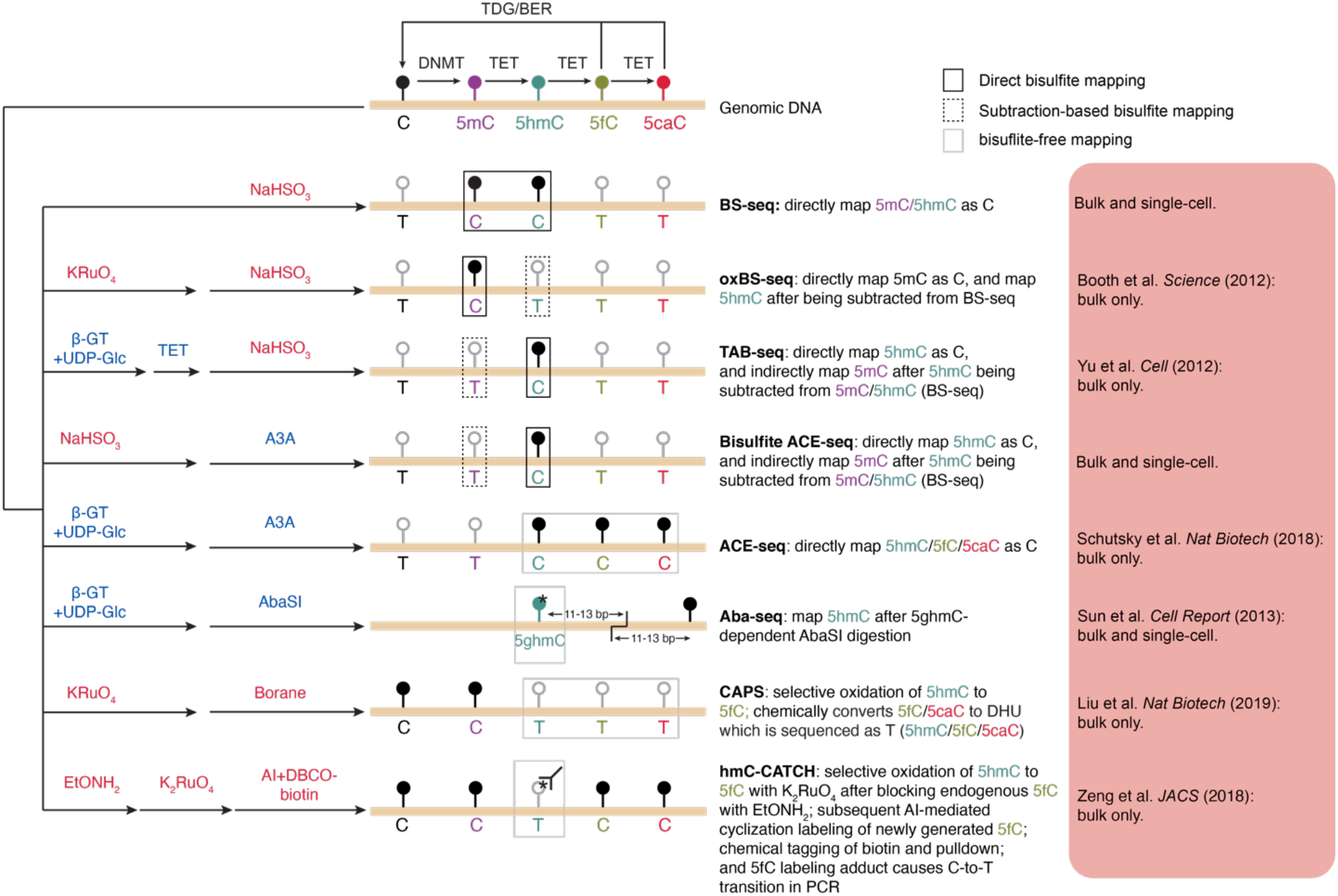
Summary of whole-genome, base-resolution 5hmC sequencing methods. A schematic comparison of epigenetic sequencing technologies used to directly (TAB-seq, Aba-seq, ACE-seq, bisulfite ACE-seq, CAPS, and hmC-CATCH) or indirectly (oxBS-seq: require subtraction from BS-seq signals) profile 5hmC through various chemical (red) and/or enzymatic (blue) reactions, followed by high-throughput sequencing. Note that 5-formylcytosine (5fC) and 5-carboxylcytosine (5caC) are of very low abundance in wild-type cells ^11^, so their contribution to 5hmC signals can be generally ignored in ACE-seq and CAPS. In AbaSI-assisted sequencing (Aba-seq)^15^, robust activity of 5ghmC-dependent AbaSI restriction endonuclease requires two cytosines on opposite strands flanking the cleavage sites, with at least one cytosine being modified for efficient cutting^36^. The absence of the second cytosine in the DNA sequence significantly reduces AbaSI activity, therefore presenting challenges for unbiased genomic coverage. The modification state of the second cytosine also influences the enzyme activity (from high to low: 5ghmC > 5hmC > 5mC > C)^15^. If both cytosines are in CG or CH contexts (accounts for 13% of all identified sites in mouse ESCs), 5hmC site cannot be assigned with certainty^15^. NaHSO_3_, sodium bisulfite; TET, Ten-eleven translocation family of DNA dioxygenase; A3A, human APOBEC3A DNA deaminase; KRuO_4_, potassium perruthenate; βGT, β-glucosyltransferase; 5ghmC, β-glucosyl-5 - hydroxymethylcytosine; UDP-Glc, UDP-Glucose; EtONH_2_, *O*-ethylhydroxylamine; K_2_RuO_4_, potassium ruthenate; AI, azido derivative of 1,3-indandione.

**Figure S2:**
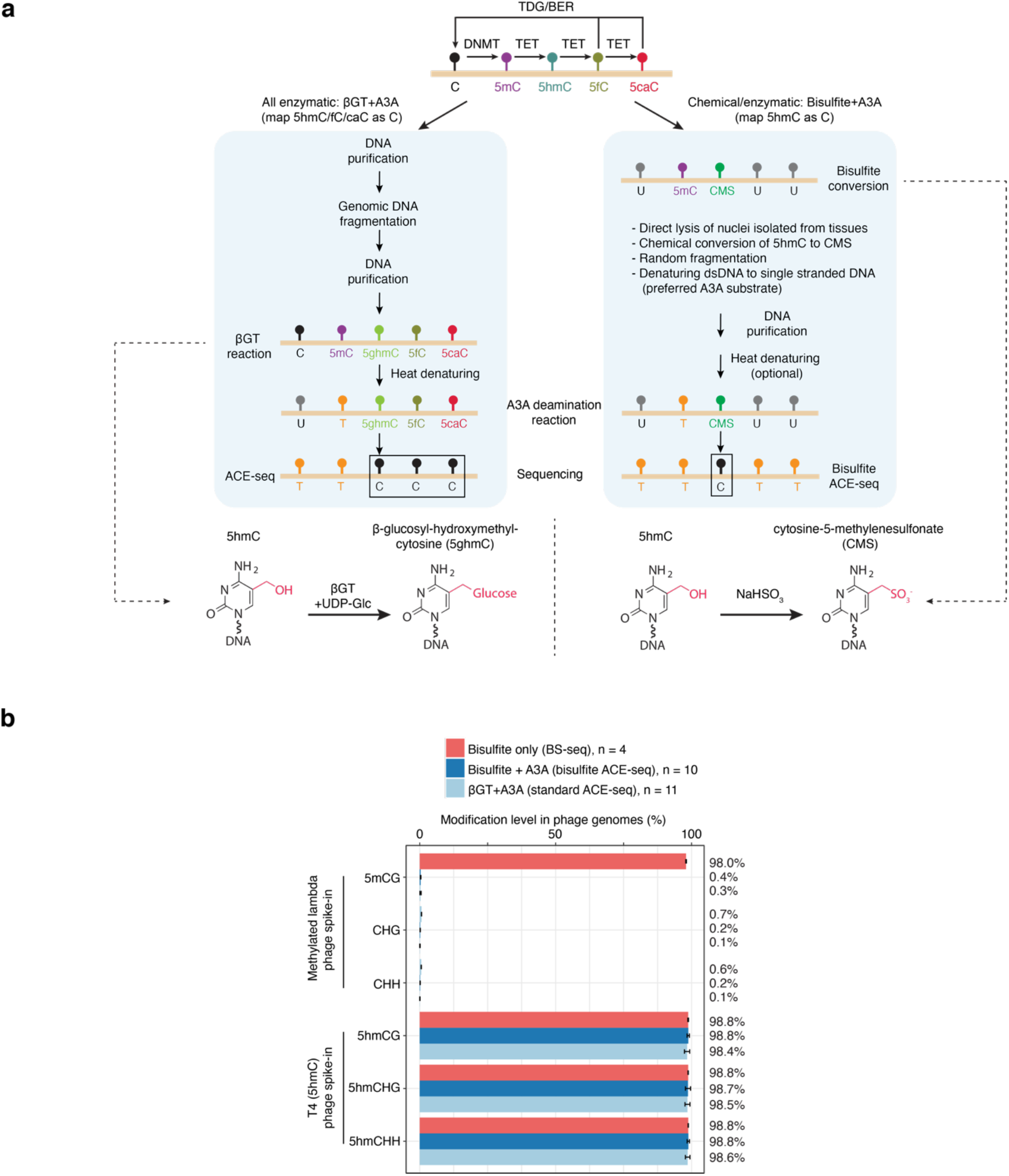
Development and validation of bisulfite ACE-seq for whole genome 5hmC analysis. **a,** Workflow comparison illustrating stepwise differences in cytosine modification status and 5hmC- protection between standard and bisulfite ACE-seq, respectively. **b,** Bar plots showing the cytosine modification levels in spike-in controls (methylated lambda and mutant T4 phage genomes) analyzed by BS-seq (red), bisulfite ACE-seq (dark blue) and standard ACE-seq (light blue). All methylated (5mCG) and unmodified cytosines (CHG/CHH) in the lambda phage genome should be deaminated in standard and bisulfite ACE-seq. Conversely, 5hmC should be protected at every cytosine position in the mutant T4 (5hmC only) phage genome for all three techniques. The mean values of multiple replicates are shown on the right. Bars and error bars represent the mean ± s.d. of results from multiple experiments (number of replicates are indicated on the top).

**Figure S3:**
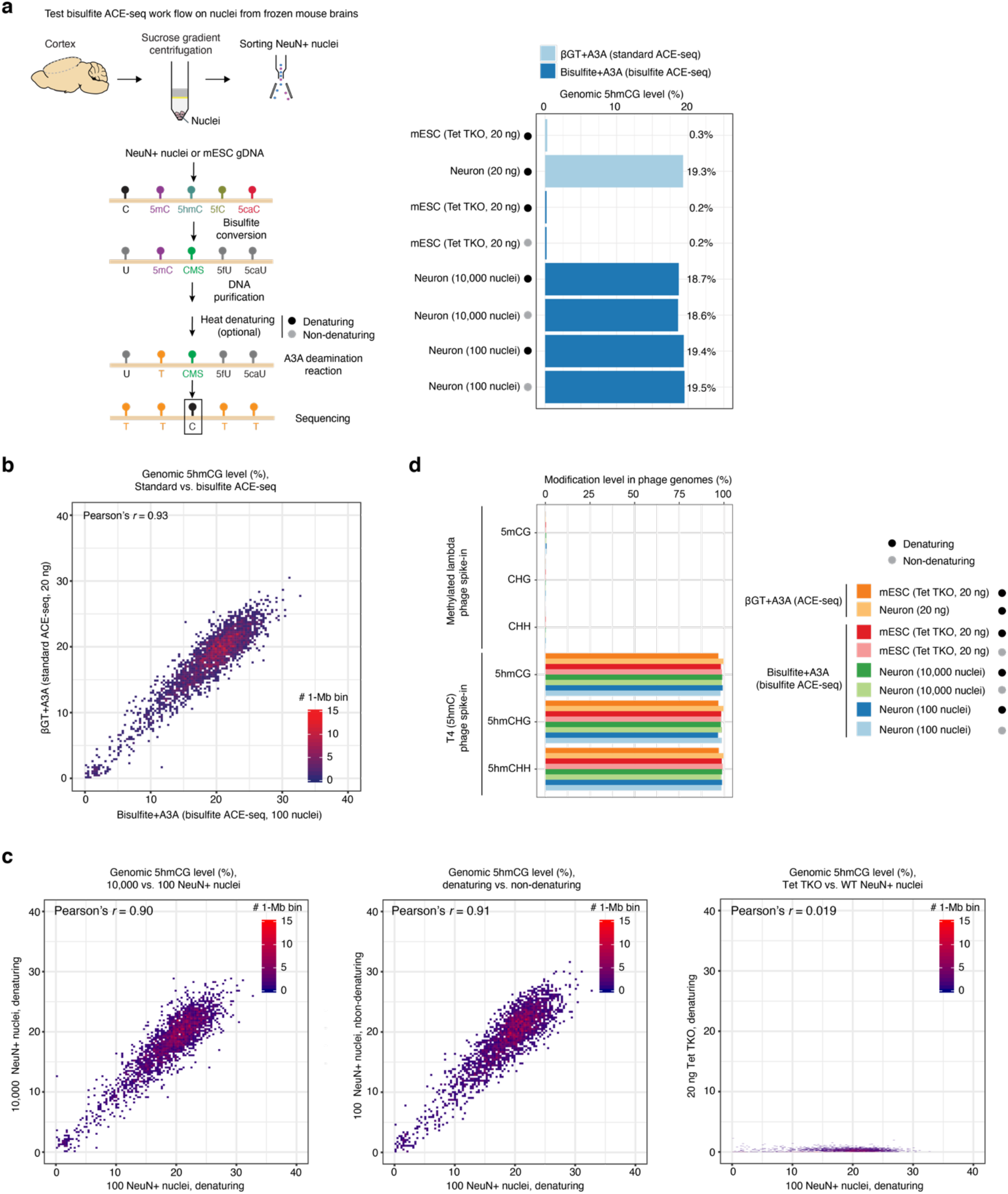
Evaluation of bisulfite ACE-seq in analysis of nuclei from cryo-preserved tissue samples. **a,** Workflow of bisulfite ACE-seq analysis of neuronal nuclei isolated from adult mouse brains (left). Bar plot in the right panel shows the global 5hmCG level in Tet1/2/3 triple KO (Tet TKO) mESCs and mouse cortical neurons analyzed by various sample preparation protocols. As a positive control, ACE-seq (light blue) was used to analyze sheared and denatured gDNA from Tet TKO mESCs or wild-type mouse cortical neurons. Bisulfite ACE-seq was applied to purified Tet TKO mESC gDNA (as a negative control) and sorted NeuN+ nuclei from mouse cortex. The black dot indicates experiments with DNA denaturation, while the grey dot indicates experiments without DNA denaturation. **b,** Correlation density plot comparing 5hmCG levels in 1-Mb bins between standard ACE-seq (20ng purified, excitatory neuronal DNA) and bisulfite ACE-seq (10,000 neuronal NeuN+ nuclei). **c,** Correlation density plot comparing global 5hmC level in 1-Mb bins between 10,000 and 100 NeuN+ nuclei with a pre-A3A denaturation step (left), between 100 NeuN+ nuclei with the pre-A3A denaturing step and without denaturing (middle), and between Tet TKO mESC DNA and 100 NeuN+ nuclei (right). **d,** Bar plots showing the cytosine modification levels in spike-in phage genomes between standard and bisulfite ACE-seq. The 5mCG and C deamination efficiency was assessed by the methylated lambda phage, and 5hmC protection was assessed by the mutant T4 (5hmC only) phage genome. Samples (same as in **a**) were indicated by colors.

**Figure S4:**
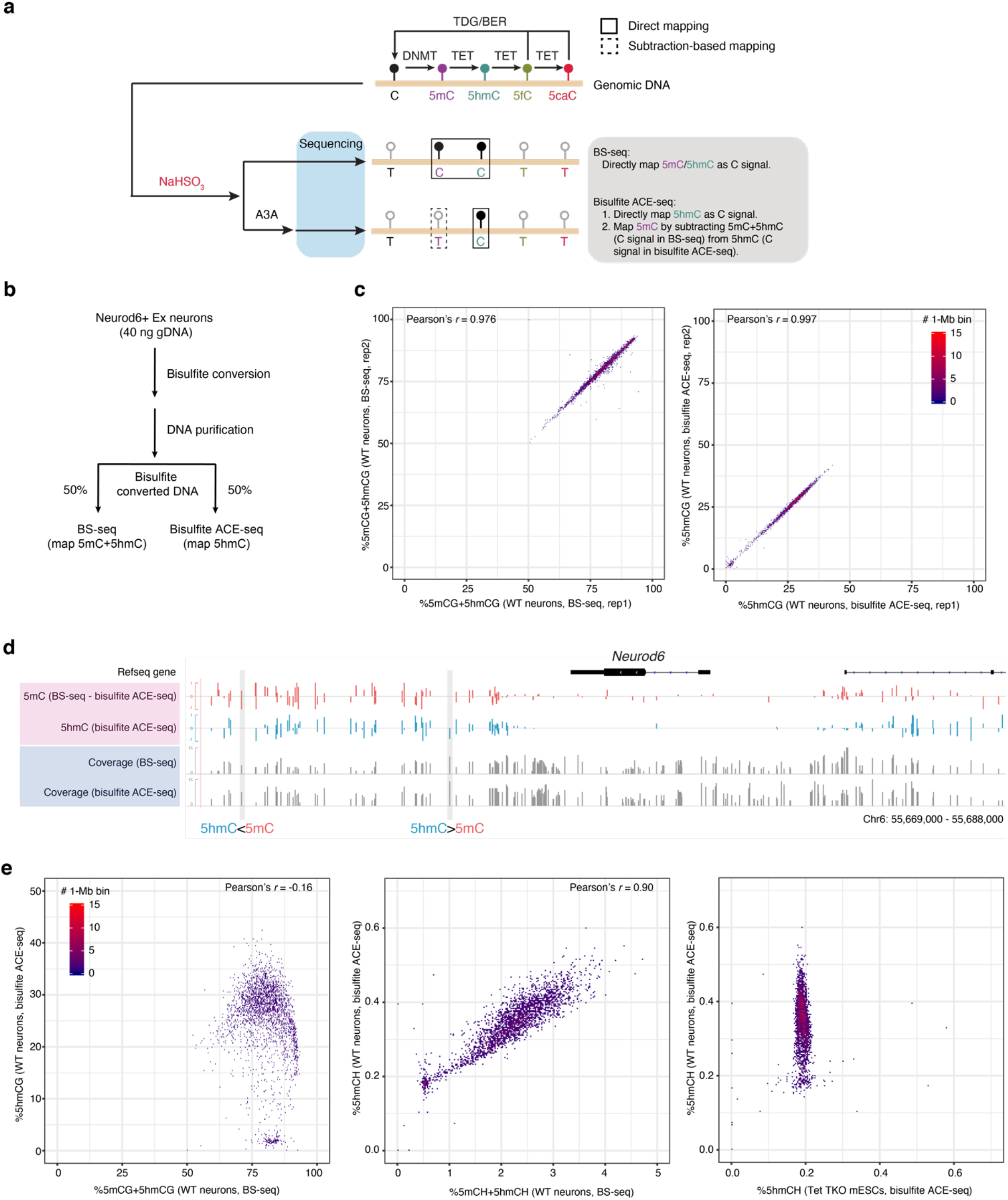
Integrated analysis of excitatory neuronal genomic DNA using paired whole genome BS-seq and bisulfite ACE-seq. **a,** The strategy of mapping both 5hmC and true 5mC by paired BS-seq and bisulfite ACE-seq. **b,** Schematic of the sample-splitting workflow for paired BS-seq and bisulfite ACE-seq on the same pool of mouse excitatory neurons. **c,** Scatterplots showing technical reproducibility of BS-seq (left, 5mCG+5hmCG) and bisulfite ACE-seq (right, 5hmCG) between two biological replicates. **d,** Genome browser tracks of 5hmCG (blue) and true 5mCG (red) signals at the *Neurod6* gene locus. The 5mC level was calculated by directly subtracting bisulfite ACE-seq signals from BS-seq. Two representative CG sites with different levels of 5mC and 5hmC were highlighted with a grey box. The sequencing coverage of the two methods are shown in grey tracks, respectively. **e,** Correlation density plot comparing genome-wide 5mCG+5hmCG (left) and 5mCH+5hmCH (middle) with 5hmCG and 5hmCH, respectively, across 1- Mb bins from WT mouse cortical neurons. The correlation density plot in the right panel compares genome-wide 5hmCH level in 1-Mb bins between WT mouse cortical neurons and Tet TKO mESCs.

**Figure S5:**
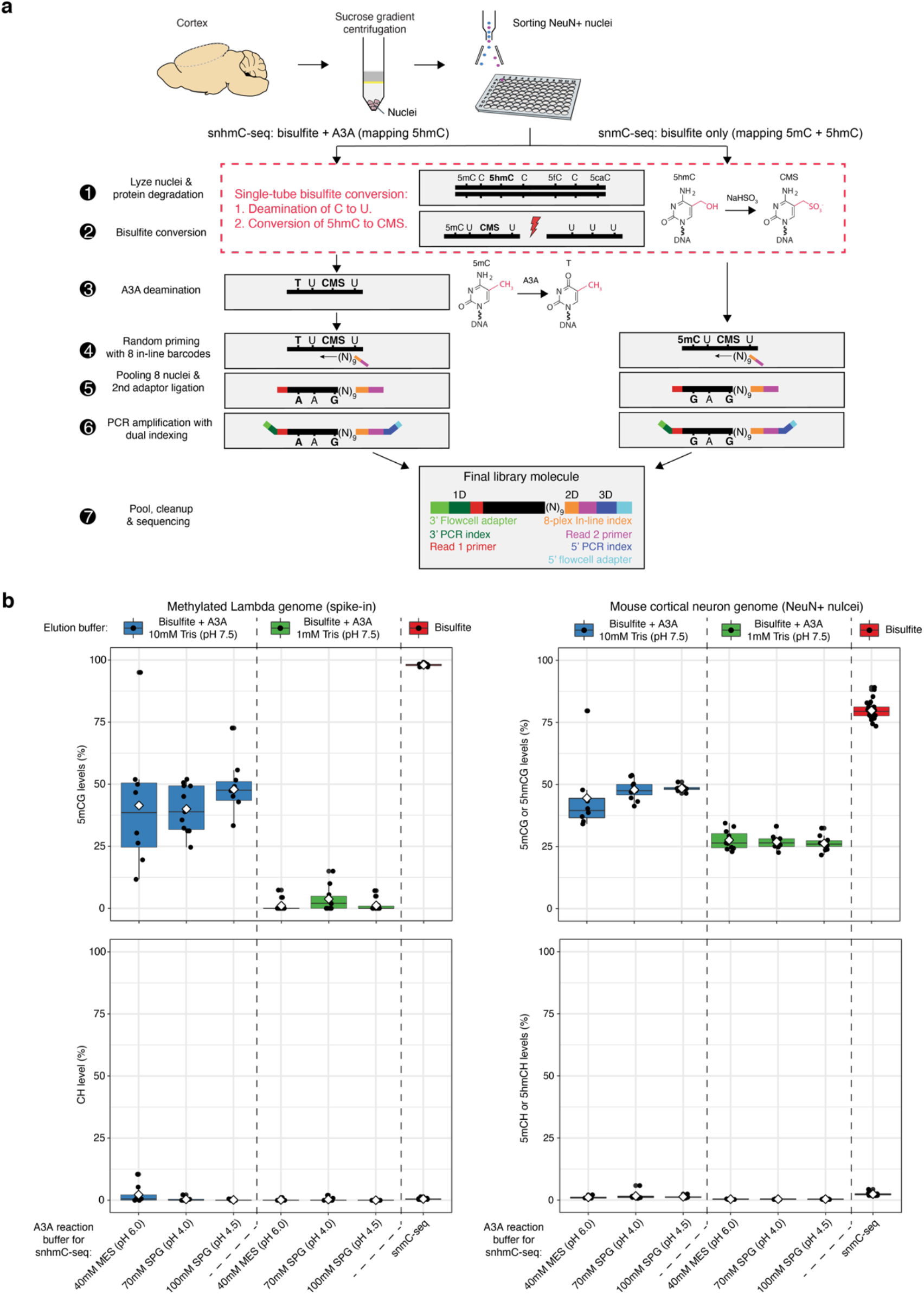
snhmC-seq workflow and buffer optimization for A3A reactions. **a,** Schematic comparing the library preparation steps between snhmC-seq and snmC-seq (Luo et al.^4^). **b,** Box plots showing the cytosine modification states within CG (top) and CH (bottom) contexts in the spike- in methylated lambda genome (left) and mouse cortical neurons (right) between different pre-A3A elution buffer and A3A reaction buffer conditions. Single-cell BS-seq (snmC-seq) was also performed in parallel as a control. Three different A3A reaction buffer conditions (based on either MES or SPG buffers) were tested. Tris-HCl elution buffer concentrations (10 mM versus 1 mM) were also evaluated to achieve optimal A3A reaction conditions in snhmC-seq assay. MES: 2-(N-morpholino)ethanesulfonic acid; SPG: a mixture of succinic acid, sodium dihydrogen phosphate, and glycine in the molar ratios 2:7:7.

**Figure S6:**
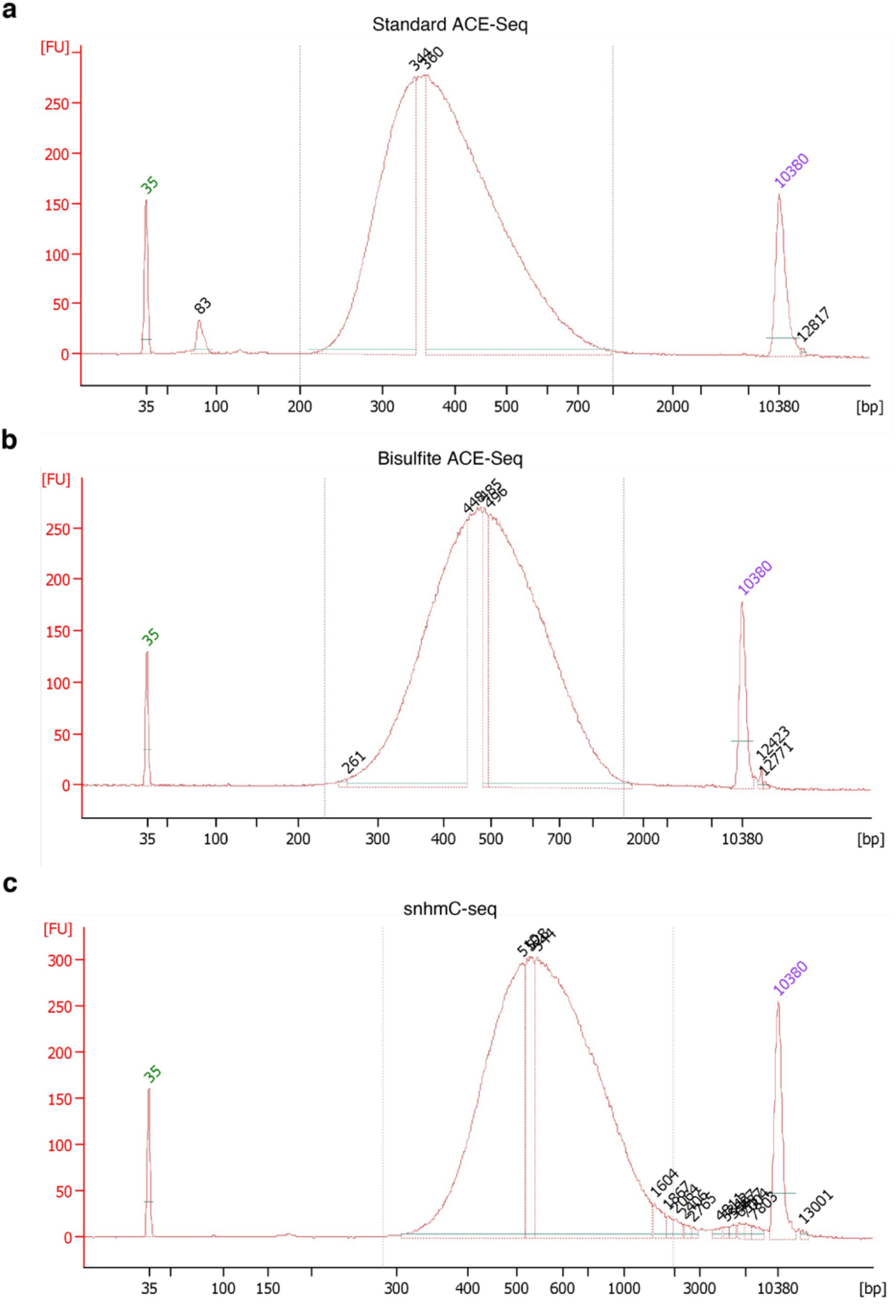
Representative bioanalyzer traces of bulk and single-cell 5hmC profiling method. Representative Bioanalyzer traces of **(a)** standard ACE-seq, **(b)** bisulfite ACE-seq, and **(c)** snhmC-seq post indexing PCR libraries. We note that standard ACE-seq libraries were initially sheared to ∼300bp via Covaris fragmentation, whereas bisulfite ACE-seq and snhmC-seq rely solely on bisulfite-mediated chemical fragmentation and random priming based extension.

**Figure S7:**
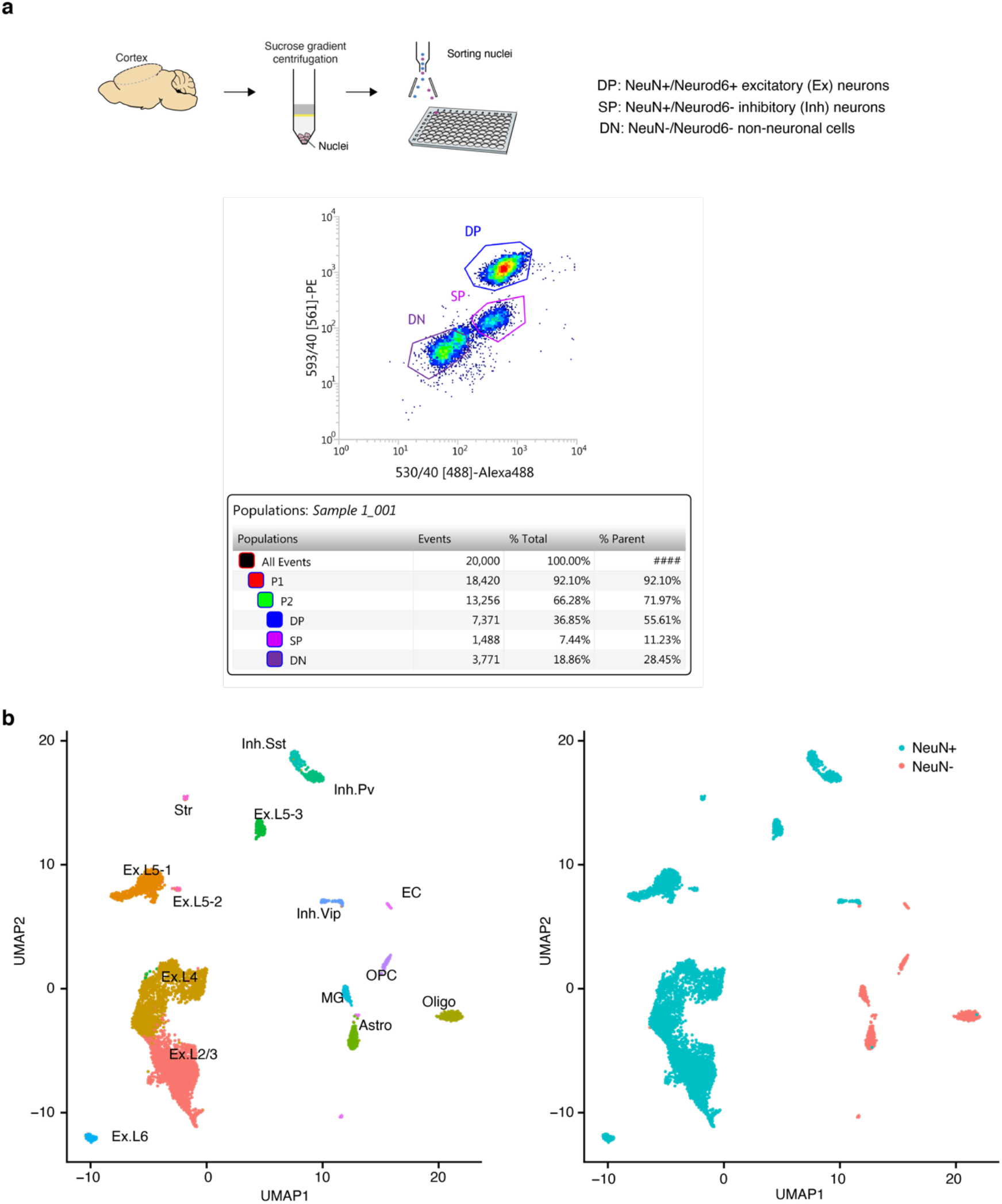
FANS of excitatory (Ex) and inhibitory (Inh) neuronal subtypes from the mouse cortex. **a,** Representative FANS results (x-axis: NeuN immunostaining signals; y-axis: Nex/Neurod6-Cre signals) for isolating excitatory neurons (double positive (DP): NeuN+/Neurod6+), inhibitory neurons (single positive (SP): NeuN+/Neurod6-), and non-neuronal cells (double negative (DN): NeuN-/Neurod6-) from the mouse cortex of *NeuroD6/NEX-Cre* mice^29^. **b,** UMAP visualization of mouse cortical nuclei colored by cellular identity defined by single-nucleus RNA sequencing analysis (left panel) or FANS analysis (right panel). Ex: excitatory neurons. Inh: inhibitory neurons; L: layer, Str: striatum; Astro, astrocytes; OPC, oligodendrocyte precursor cells; Oligo, oligodendrocytes; MG, microglia; EC, endothelial cells.

**Figure S8:**
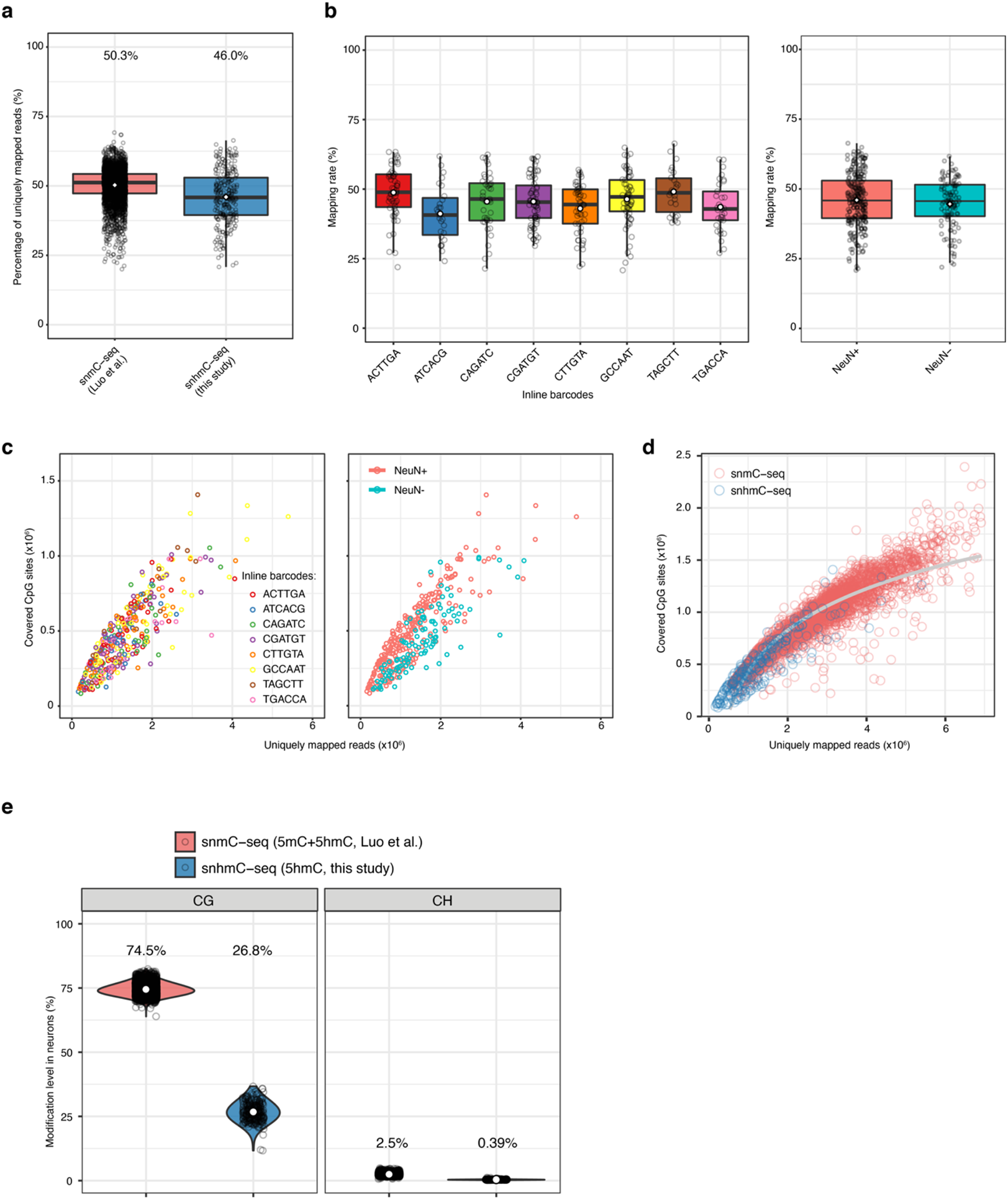
Benchmarking snhmC-seq performance. **a,** Boxplots showing the percentage of uniquely mapped reads (over total reads) in snhmC-seq and snmC-seq (Luo et al.^4^). Open white circles indicate the mean value. See ‘Data visualization’ in the Methods for definitions of boxplot elements. **b,** Boxplots showing the mapping rates across 8-plex inline barcodes (left panel) or mapping rates associated with neuronal (NeuN+) and non-neuronal (NeuN-) nuclei (right panel). Open circles indicate the mean value. **c,** Scatterplots showing the covered CpG sites per nucleus as a function of uniquely mapped reads per nucleus for 8-plex inline barcodes (left panel) and different cell populations sorted via FANS (right panel). **d,** Scatterplot comparing the covered CpG sites per nucleus as a function of uniquely mapped reads per nucleus between snmC-seq (red) and snhmC-seq (blue). The grey line indicates the fitted curve. **e,** Violin plots showing the global cytosine modification levels within CG (left panel) or CH (right panel) contexts detected in snmC-seq and snhmC-seq. Open circles indicate the mean value. See ‘Data visualization’ in the Methods for definitions of violin plot elements.

**Figure S9:**
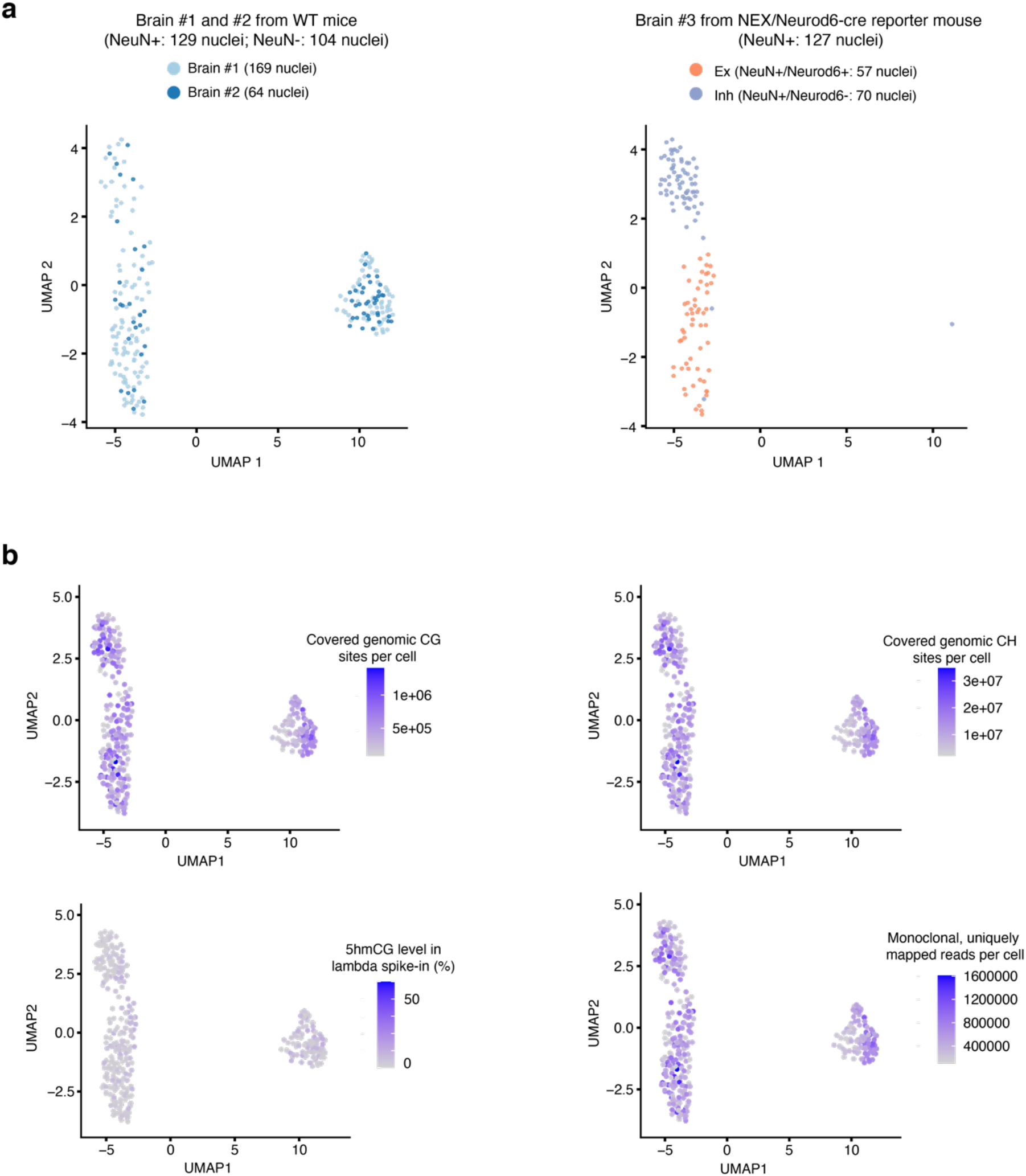
Quality control of unbiased clustering results of mouse cortical nuclei. **a,** UMAP visualization (same as Fig. 1e) of 233 sorted cortical nuclei from wild-type mouse (brain #1 and 2) colored by biological replicates (left panel) and of 127 sorted mouse cortical nuclei from Nex-Cre transgenic mouse (brain #3) colored by cell-types defined by FANS analysis (right panel). **b,** UMAP visualization (same as Fig. 1e) showing covered CG sites, number of covered CH sites, global 5hmCG level in the lambda spike-in, and number of uniquely mapped reads per nucleus.

**Figure S10:**
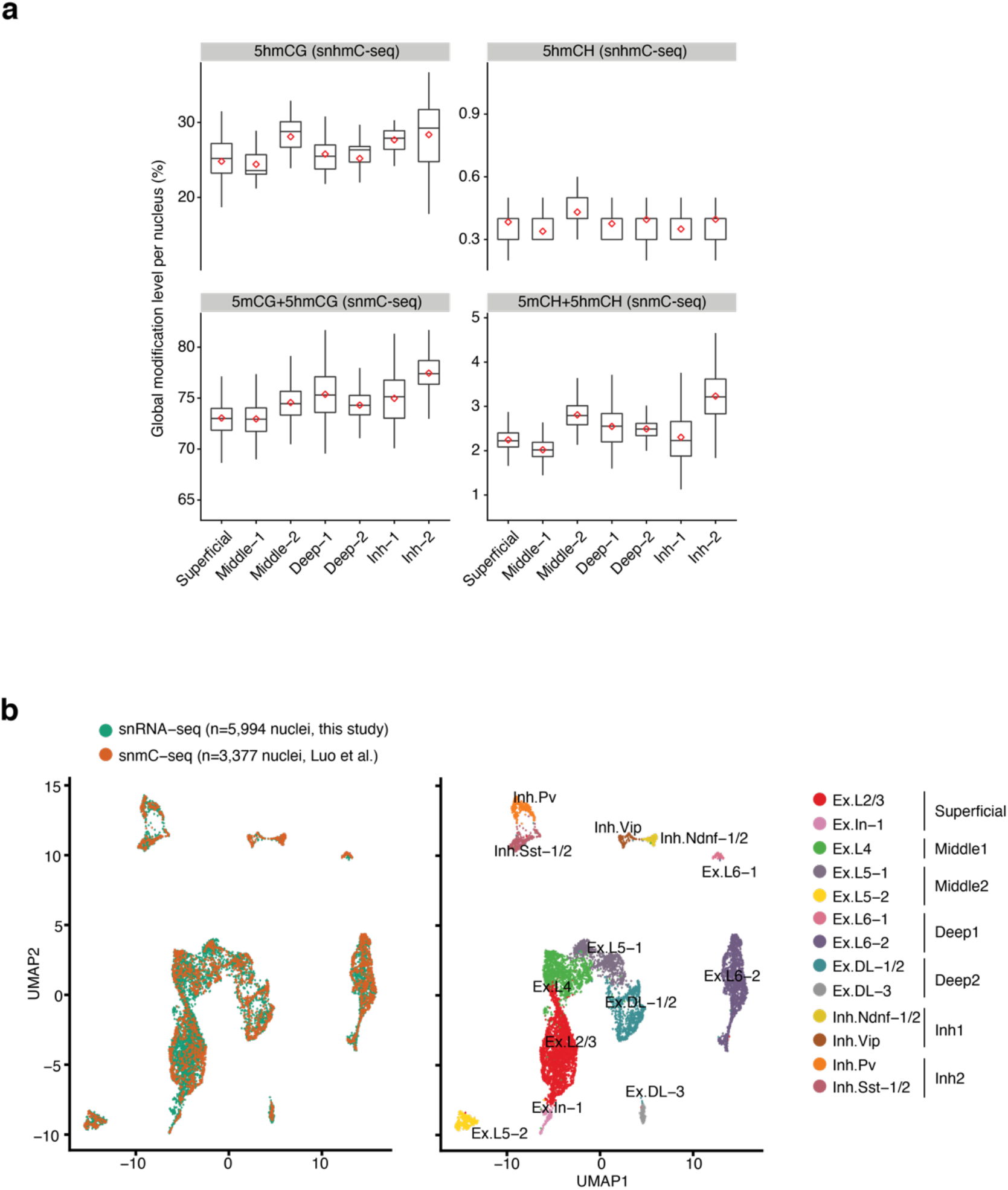
Joint analysis of snhmC-seq, snmC-seq and sNucDrop-seq. **a,** Box plots showing global CG and CH modification levels across 7 major mouse cortical neuronal subtypes. Red dots indicate the average level of each modification. See ‘Data visualization’ in the Methods for definitions of box plot elements. **b,** UMAP visualization of 9,371 mouse cortical NeuN+ nuclei from joint analysis of sNucDrop-seq (in this study) and snmC-seq (Luo et al.^4^), colored by different methods (left panel) or cluster assignment (right panel). Ex: excitatory neuron; Inh: inhibitory neuron; L: layer; DL: deep layer.

## METHODS

### Cell lines and culture conditions

For bisulfite ACE-seq validation experiments: *Tet*1/2/3 triple-knock out (Tet TKO) mESCs (J1) were initially cultured in presence of Mitomycin C inactivated mouse embryonic fibroblasts on 0.1% gelatin-coated (Millipore, ES-006-B) 6-well plates in Dulbecco’s Modified Eagle’s Medium (DMEM) (Gibco, 11965084) supplemented with 15% fetal bovine serum (Gibco, 16000044), 0.1 mM nonessential amino acid (Gibco, 11140050), 1 mM sodium pyruvate (Gibco, 11360070), 2 mM L-glutamine (Gibco, 25030081), 50 μM 2-mercaptoethanol (Gibco, 31350010), 1 μM MEK inhibitor PD0325901 (Axon Med Chem, Axon 2128) and 3 μM GSK3 inhibitor CHIR99021 (Axon Med Chem, Axon 2128), and 1,000 U/mL LIF (Gemini Bio-Products, 400-495-7). Generation and genotyping of *Tet*-TKO mESCs were previously described^24^.

### Animals

Experiments were conducted in accordance with the ethical guidelines of the National Institutes of Health and with the approval of the Institutional Animal Care and Use Committee of the University of Pennsylvania. Mice were group-housed in cages of three to five in a 12-hr light/dark cycle with food and water provided ad libitum. All mice used for experiments were naive to behavioral assays and other procedures. For all bulk-nucleus and single-nucleus experiments, 6- to 10-week-old, wild type, male mice in a pure C57BL/6J background were used. For prospectively isolating inhibitory and excitatory neuronal subtypes to benchmark snhmC-seq assay, 9-week-old transgenic mice (*Mecp2^Tavi/y^*, *Nex^Cre/+^*, *Rosa26^BirA/BirA^*) from a C57BL/6J background were used ^29^.

### Sample preparation and nuclear isolation

Mouse brains (postnatal 6-10 weeks) were rapidly resected on ice. Cortices were flash frozen in liquid nitrogen for 2min and subsequently kept at −80°C before nuclear isolation. Nuclei were isolated and purified as previously described with some modifications^3^. Briefly, 14mL sucrose cushion (1.8M sucrose (Sigma-Aldrich, RNase and DNase free, ultra-pure grade), 10mM Tris-HCl pH 8.0, 3mM MgAc_2_ (Sigma-Aldrich), protease inhibitor cocktail (Roche)) was added to the bottom of centrifuge tubes (Beckman Coulter). Using a glass homogenizer (Wheaton), a frozen mouse cortex sample was subject to dounce homogenization (19 times with loose pestle followed by 4 times with tight pestle) in 12mL homogenization buffer (0.32M sucrose, 5mM CaCl_2_ (Sigma-Aldrich), 3mM MgAc_2_, 10mM Tris-HCl pH 8.0, 0.1% Triton X-100 (Sigma-Aldrich), 0.1mM EDTA (Invitrogen), protease inhibitor cocktail). Homogenates (∼12mL) were carefully layered onto the sucrose cushion in the centrifuge tubes, and 10mL homogenization buffer was added atop of the homogenates. The tubes were then balanced and centrifuged in a Beckman Coulter L7-65 Ultracentrifuge at 25,000rpm at 4°C for 2hr using a Beckman Coulter SW28 swing bucket rotor (Beckman Coulter). The supernatant was carefully removed via aspiration. 1mL chilled DPBS was added to resuspend the nuclear pellet, and nuclei were subsequently transferred to a 1.5-mL tube. Nuclei were pelleted at 5,000 rpm for 10 min at 4C, and then resuspended in 0.01% BSA (Sigma-Aldrich) in DPBS. After resuspension, nuclei were filtered through a 40-mm cell strainer (Fisher Scientific), visually inspected for morphology and quality assurance, and counted using a Fuchs-Rosenthal counting chamber before FANS.

### Flow cytometry-based sorting of NeuN+/− nuclei

After filtering, nuclei were resuspended in blocking buffer (1x PBS, 0.5% BSA) and pelleted using a tabletop centrifuge at 5,000 RCF at 4°C for 15min. Nuclei were resuspended in 1.4mL blocking buffer (1x PBS, 0.5% BSA). 1mL of this cell suspension was transferred to a separate 1.5mL Eppendorf tube, and the remaining 0.4mL were split between two additional 1.5mL Eppendorf tubes (to later serve as FANS controls). 800µL blocking buffer was added to one of the two 1.5mL Eppendorf tubes containing 0.2mL of the counted nuclear suspension, and 1mL to the other to supplement the volume of these reactions for future antibody incubation. During this time, 1.2µL NeuN+ primary antibody (Millipore, MAB377) and 1µL Alexa488 secondary antibody (Invitrogen) were incubated at room temperature for 5min in 150µL 1x PBS and 50µL blocking buffer: primary + secondary antibody solution. This same mixture was incubated, minus the primary antibody, to serve as a negative FANS control (secondary-only control). The primary + secondary antibody solution was added to the undiluted nuclear suspension, and the secondary-only control was added to the 1mL diluted nuclear suspension. Both of these tubes were covered with aluminum foil and incubated with rotation at 4°C for 25min. Subsequently, 1.2µL 100mg/µL DAPI (Sigma-Aldrich) was added to all three nuclear suspensions. The tubes were covered with aluminum foil, and further incubated at 4°C with rotation for 5min. All three suspensions were pelleted using a tabletop centrifuge at 5,000 RCF at 4°C for 15min. The supernatants were removed and 1mL blocking buffer was added to resuspend each pellet. All three suspensions were washed at 4°C with rotation for 20min. All three suspensions were pelleted using a tabletop centrifuge at 5,000 RCF at 4°C for 15min. Each pellet was resuspended to a final concentration of 1×10^6^nuclei/mL and sorted with a BD Biosciences Influx cell sorter at the University of Pennsylvania Flow Cytometry and Cell Sorting Facility. DAPI-only and secondary-only control samples were used to calibrate the sorter, and NeuN+/− nuclear populations were either sorted in bulk (into 1.5mL Eppendorf tubes) or single-nuclei (in 96-well PCR plates provided in the Zymo EZ-96 DNA Methylation-Direct™ Kit loaded with 4µL Proteinase K digestion buffer (1µL M-Digestion Buffer, 0.1µL 20µg/µL Proteinase K, and 0.9µL water).

### Flow cytometry-based sorting of excitatory and inhibitory neurons using the *Neurod6/NEX*-Cre driven transgenic reporter mouse line

A transgenic reporter mouse line was generated as previously described^29^. Isolated nuclei from prefrontal cortex were resuspended in 1.5mL blocking buffer (1× PBS, 0.5% BSA (Sigma A4503), Roche Complete Protease Inhibitor without EDTA) and blocked for 20 min at 4 °C with rotation. After blocking, nuclei were incubated with an Alexa 488-conjugated anti-NeuN antibody (1:1,000, Millipore MAB377X) and Streptavidin-Phycoerythrin (PE) (1:200, BioLegend 405203) for 30 min at 4 °C with rotation. After a 5 min incubation with DAPI (1:1000), nuclei were washed for an additional 30 min in 1mL blocking buffer. Nuclei were then pelleted, resuspended in 1mL FACS buffer (1× PBS, 1% BSA (Sigma A4503), Roche Complete Protease Inhibitor without EDTA), and passed through a 35 µm strainer (Corning #352235) in preparation for flow cytometry. FACS was performed using a BD Biosciences Influx cell sorter at the University of Pennsylvania Flow Cytometry and Cell Sorting Facility. Singlet nuclei were identified using DAPI fluorescence, and Alexa 488 and PE fluorescence were used to isolate Streptavidin-/NeuN- (non-neuronal, double negative in **Fig S7a**), Streptavidin-/NeuN+ (*Nex*- negative neuronal nuclei, Inh, single positive in **Fig S7a**), and Streptavidin+/NeuN+ (*Nex*-positive neuronal nuclei, Ex, double positive in **Fig S7a**) populations.

### Preparation of phage spike-in controls

Phage spike-ins (used in standard/bisulfite ACE-seq and snmC/snhmC-seq assays) were prepared as previously described^1^, with minor modifications. Briefly, lambda phage DNA was enzymatically methylated at CG sites by incubating with the CG methyltransferase M.SssI (NEB M0226M) and S-adenosylmethionine (SAM) at 37°C as previously reported^23^. After 2h, additional enzyme and SAM were added to the reaction and incubation continued for another 4h, the enzyme was deactivated by incubation at 65°C for 20min, and DNA was bead purified with 1.6x homebrew solid phase reversible immobilization (SPRI) beads (1mL Sera-Mag SpeedBeads (GE Healthcare), 9g PEG 8000 (Sigma), 10mL 5M NaCl, 500µL 1M Tris-HCl pH 8.0 (Invitrogen), 100µL 0.5M EDTA pH 8.0 (Invitrogen)). The methylated phage DNA was combined in equal amounts with mutant T4 (5hmC only) phage DNA (Schutsky et al. 2018). Unsheared phage DNA is added in a proportion of 1:200 for bulk assays (standard and bisulfite ACE-seq) and 1:100 for single cell assays (snmC-seq and snhmC-seq).

### APOBEC3A quality control for standard and bisulfite ACE-seq reactions

Wild-type human APOBEC3A (A3A) was cloned, expressed, and purified from BL21 (DE3) cells with a Trigger Factor (TF) chaperone, as previously described^2^. Each batch of purified enzyme must undergo quality control measures to ensure normalized batch-specific reaction conditions. To validate optimal enzyme concentration, phage-only samples were analyzed with 10ng each of pooled methylated phage and mutant T4 (5hmC only) gDNA. Each batch of A3A enzymes were tested on spike-in controls at the measured concentration plus seven subsequent 1:2 serial dilutions and methylated phage deamination efficiency, as well as 5hmC protection efficiency were validated by sequencing a pooled 4nM library on the Illumina MiSeq using the MiSeq Reagent Nano Kit v2 (300-cycles).

### APOBEC3A reaction buffer optimization for low-input bisulfite ACE-seq and snhmC-seq

Due to the difference in the proportion of reaction components between standard and bisulfite ACE-seq, buffer optimization was performed to ensure the high efficiency of the APOBEC3A deamination reaction in the single-cell 5hmC mapping workflow **(Fig. S5b)**. The ACE-seq reaction supports up to 39% of the A3A incubation reaction volume to consist of eluted DNA, whereas bisulfite ACE-seq and snhmC-seq require 60% of this final reaction volume to consist of DNA eluent. As a result, buffer conditions for this step were specifically optimized to maintain pH 6.0 reaction conditions for this incubation step. Two strategies were used to achieve optimal reaction condition: 1) reducing the concentration of the Tris-HCl pH 7.5 elution buffer from 10mM to 1mM, post bisulfite treatment; 2) Utilizing higher concentration of standard ACE-seq buffer in the final reaction (40 mM MES pH 6.0)^4^ and an alternative reaction solution, SPG buffer ^37^ (a mixture of succinic acid, sodium dihydrogen phosphate, and glycine in the molar ratios 2:7:7) that buffers at a wider pH range **(Fig. S5b)**.

### Standard and bisulfite ACE-seq comparison on phage controls and mammalian genomic DNA

Standard ACE-seq was performed as previously described^17^. For all mammalian genomic DNA samples, a total of 20ng of sheared gDNA (∼300bp) was analyzed, containing the sheared methylated lambda phage and mutant T4 phage (5hmC only) spike-in controls (0.5% total by mass). In a total volume of 5uL, the gDNA mixture was glucosylated using UDP-glucose and T4 ꞵ-glucosyltransferase (T4-ꞵGT, NEB M0357S) at 37°C for 1h. 1µL DMSO was added, and the sample was denatured at 95°C for 5min and snap cooled by transfer to a PCR tube rack pre-incubated at −80°C. Before thawing, reaction buffer was overlaid to a final concentration of 20mM MES pH 6.0 + 0.1% Tween, and A3A was added to a final concentration of 5μM in a total volume of 10μL. The deamination reactions were incubated under linear ramping temperature conditions from 4–50°C over 2h. After deamination, the samples were prepared for Illumina sequencing using the Accel Methyl-NGS kit (Swift Biosciences, 33096). Bisulfite ACE-seq was performed on 20ng unsheared DNA containing unsheared methylated lambda phage and mutant T4 phage (5hmC only) spike-in controls (0.5% total by mass). Bisulfite conversion and purification was carried out using EZ DNA Methylation-Direct™ Kit (Zymo Research D5020), following the product manual. Bisulfite reactions were eluted with 10µL 0.5mM Tris-HCl pH 7.5. To 9µL of this reaction, 1.5µL 40mM MES pH 6.0 + 0.1% Tween and 1.5µL DMSO were added. The sample was denatured (optional for bisulfite ACE-seq) at 95°C for 1min and snap cooled by transfer to a PCR tube rack pre-incubated at −80°C. Before thawing, 1.5µL 40mM MES pH 6.0 + 0.1% Tween and 1.5µL 50μM A3A were added to each reaction to a final volume of 15µL. The deamination reactions were incubated at 37°C for 2h, purified with 1.6x homebrew SPRI beads, and prepared for Illumina sequencing using the adapted snmC-seq strategy (see below)^4^.

### Library preparation of low-input bisulfite ACE-seq and snhmC-seq

All steps of library preparation were performed in an AirClean^®^600 PCR Workstation to minimize environmental DNA contamination. Bisulfite conversion of bulk nuclei for bisulfite ACE-seq was carried out using Zymo EZ DNA Methylation- Direct™ Kit (D5020), following the product manual. Bisulfite conversion of single nuclei for snhmC-seq was carried out using Zymo EZ-96 DNA Methylation-Direct™ Kit (Deep Well Format, D5023), following the product manual with proportionally reduced reaction volume. PCR tubes containing FANS-isolated nuclei, or 96-well plates containing FANS-isolated single nuclei, were heated at 50°C for 20min. 130μL (for bulk) or 32.5μL (for single nuclei) CT Conversion Reagent was added to each well, followed by pipetting up and down to mix. Plates were treated with the following program using a thermocycler: 98°C for 8min, 64°C for 3.5h, and 4°C forever.

Zymo-Spin™ columns or Zymo-Spin™ I-96 Binding Plates were preloaded with either 600μL or 250μL M-binding buffer, respectively. Bisulfite conversion reagents were transferred from their reaction vial into their respective spin columns or plates. From this point, both bulk and single-cell samples were further processed according to their respective kit instructions, except 0.5mM Tris-HCl pH 7.5 was used as the elution buffer. To the 9μL eluent retained from each reaction, and to this, 1.5µL 200mM MES pH 6.0 + 0.1% Tween and 1.5µL DMSO were added. The samples were denatured (optional for bisulfite ACE-seq) at 95°C for 1min and snap cooled by transfer to a PCR tube rack pre-incubated at −80°C (for bulk samples) or transfer to an isopropanol dry ice bath and then to the −80°C pre-incubated PCR tube rack (for 96-well plates containing single nuclei). Before thawing, 1.5µL 200 mM MES pH 6.0 + 0.1% Tween- 20 and 1.5µL 5μM A3A were added to each reaction to a final volume of 15µL (for a final concentration of 500nM/µL A3A per reaction). The deamination reactions were incubated at 37°C for 2h, purified with 1.6x homebrew SPRI beads, eluted in 9µL Low EDTA TE buffer (Swift Biosciences) and prepared for Illumina sequencing using the adapted snmC-seq library preparation workflow as previously outlined^5, 6^.

Briefly, 1µL of an assigned random primer (P5L_AD002_H, P5L_AD006_H, P5L_AD008_H, P5L_AD010_H, P5L_AD001_H, P5L_AD004_H, P5L_AD007_H, or P5L_AD012_H) was added to each reaction to allow for subsequent multiplexing (8-plex) reactions during downstream library preparation.

#### Random primers with 8-plex inline barcodes

(HPLC purified) were ordered from Integrated DNA Technologies (IDT):

P5L_AD002_H:

/5SpC3/TTCCCTACACGACGCTCTTCCGATCT<u>CGATGT</u>(H1:33340033)(H1)(H1)(H1)(H1)(H1)(H1)(H1)(H1)

P5L_AD006_H:

/5SpC3/TTCCCTACACGACGCTCTTCCGATCT<u>GCCAAT</u>(H1:33340033)(H1)(H1)(H1)(H1)(H1)(H1)(H1)(H1)

P5L_AD008_H:

/5SpC3/TTCCCTACACGACGCTCTTCCGATCT<u>ACTTGA</u>(H1:33340033)(H1)(H1)(H1)(H1)(H1)(H1)(H1)(H1)

P5L_AD010_H:

/5SpC3/TTCCCTACACGACGCTCTTCCGATCT<u>TAGCTT</u>(H1:33340033)(H1)(H1)(H1)(H1)(H1)(H1)(H1)(H1)

P5L_AD001_H:

/5SpC3/TTCCCTACACGACGCTCTTCCGATCT<u>ATCACG</u>(H1:33340033)(H1)(H1)(H1)(H1)(H1)(H1)(H1)(H1)

P5L_AD004_H:

/5SpC3/TTCCCTACACGACGCTCTTCCGATCT<u>TGACCA</u>(H1:33340033)(H1)(H1)(H1)(H1)(H1)(H1)(H1)(H1)

P5L_AD007_H:

/5SpC3/TTCCCTACACGACGCTCTTCCGATCT<u>CAGATC</u>(H1:33340033)(H1)(H1)(H1)(H1)(H1)(H1)(H1)(H1)

P5L_AD012_H:

/5SpC3/TTCCCTACACGACGCTCTTCCGATCT<u>CTTGTA</u>(H1:33340033)(H1)(H1)(H1)(H1)(H1)(H1)(H1)(H1)

Reactions or plates were heated at 95°C using a thermocycler for 3min to denature and were immediately chilled on ice for 2min. 10µL enzyme mix (2µL Blue Buffer (Enzymatics B0110), 1µL 10mM dNTP (NEB N0447L), 1µL Klenow 3’->5’ exo- (50U/µL, Enzymatics P7010-HC-L), and 6µL water) was added to each well and reactions were mixed by vortexing. Plates or reactions were treated with the following program using a thermocycler: 4°C for 5min, ramp up to 25°C at 0.1°C/sec, 25°C for 5min, ramp up to 37°C at 0.1°C/sec, 37°C for 60min, 4°C forever. Following this, 2μL Exonuclease 1 (20U/μL, Enzymatics X8010L) and 1μL Shrimp Alkaline Phosphatase (rSAP) (1U/μL, NEB M0371L) was added to each reaction followed by vortexing and incubation in a thermocycler at 37°C for 30min followed by 4°C forever.

18.4μL (0.8x) homebrew SPRI beads were added to each reaction; Up to 8 sample/bead mixtures can be pooled at this stage to allow for multiplexing reactions. The sample/bead mixtures were incubated at room temperature for 5min before being placed on a 96-well magnetic separator (DynaMag™-96 Side Magnet, ThermoFisher 12331D). Supernatant was removed from each well, followed by three rounds of washing with 180μL 80% ethanol. Beads were air dried at room temperature and 10μL Low EDTA TE buffer was added to each well to fully resuspend the beads. Eluted samples were then transferred to either new PCR strips (for bulk samples) or 96-well plates (for single-cell samples).

The reactions were denatured in a thermocycler at 95°C for 3min and subsequently chilled on ice for 2min. 10.5μL Adaptase master mix (2μL Buffer G1, 2μL Reagent G2, 1.25μL Reagent G3, 0.5μL Enzyme G4, 0.5μL Enzyme G5, and 4.25μL Low EDTA TE buffer; Accel-NGS Adaptase Module for Single Cell Methyl-Seq Library Preparation, Swift Biosciences 33096) was added to each reaction, followed by vortexing. Reactions were incubated in a thermocycler at 37°C for 30min then 4°C forever. Subsequently, 30μL PCR mix (25μL KAPA HiFi HotStart ReadyMix, KAPA BIOSYSTEMS KK2602, 1μL 30μM P5 indexing primer, and 5μL 10μM P7 indexing primer) were added to each well, followed by mixing with vortexing.

#### P5 indexing PCR primers

P5L_D501:

AATGATACGGCGACCACCGAGATCTACAC<u>TATAGCCT</u>ACACTCTTTCCCTACACGACGCTCT

P5L_D502:

AATGATACGGCGACCACCGAGATCTACAC<u>ATAGAGGC</u>ACACTCTTTCCCTACACGACGCTCT

P5L_D503:

AATGATACGGCGACCACCGAGATCTACAC<u>CCTATCCT</u>ACACTCTTTCCCTACACGACGCTCT

P5L_D504:

AATGATACGGCGACCACCGAGATCTACAC<u>GGCTCTGA</u>ACACTCTTTCCCTACACGACGCTCT

P5L_D505:

AATGATACGGCGACCACCGAGATCTACAC<u>AGGCGAAG</u>ACACTCTTTCCCTACACGACGCTCT

P5L_D506:

AATGATACGGCGACCACCGAGATCTACAC<u>TAATCTTA</u>ACACTCTTTCCCTACACGACGCTCT

P5L_D507:

AATGATACGGCGACCACCGAGATCTACAC<u>CAGGACGT</u>ACACTCTTTCCCTACACGACGCTCT

P5L_D508:

4AATGATACGGCGACCACCGAGATCTACAC<u>GTACTGAC</u>ACACTCTTTCCCTACACGACGCTCT

#### P7 indexing PCR primers

P7L_D701:

CAAGCAGAAGACGGCATACGAGAT<u>CGAGTAAT</u>GTGACTGGAGTTCAGACGTGTGCTCTT

P7L_D702:

CAAGCAGAAGACGGCATACGAGAT<u>TCTCCGGA</u>GTGACTGGAGTTCAGACGTGTGCTCTT

P7L_D703:

CAAGCAGAAGACGGCATACGAGAT<u>AATGAGCG</u>GTGACTGGAGTTCAGACGTGTGCTCTT

P7L_D704:

CAAGCAGAAGACGGCATACGAGAT<u>GGAATCTC</u>GTGACTGGAGTTCAGACGTGTGCTCTT

P7L_D705:

CAAGCAGAAGACGGCATACGAGAT<u>TTCTGAAT</u>GTGACTGGAGTTCAGACGTGTGCTCTT

P7L_D706:

CAAGCAGAAGACGGCATACGAGAT<u>ACGAATTC</u>GTGACTGGAGTTCAGACGTGTGCTCTT

P7L_D707:

CAAGCAGAAGACGGCATACGAGAT<u>AGCTTCAG</u>GTGACTGGAGTTCAGACGTGTGCTCTT

P7L_D708:

CAAGCAGAAGACGGCATACGAGAT<u>GCGCATTA</u>GTGACTGGAGTTCAGACGTGTGCTCTT

P7L_D709:

CAAGCAGAAGACGGCATACGAGAT<u>CATAGCCG</u>GTGACTGGAGTTCAGACGTGTGCTCTT

P7L_D710:

CAAGCAGAAGACGGCATACGAGAT<u>TTCGCGGA</u>GTGACTGGAGTTCAGACGTGTGCTCTT

P7L_D711:

CAAGCAGAAGACGGCATACGAGAT<u>GCGCGAGA</u>GTGACTGGAGTTCAGACGTGTGCTCTT

P7L_D712:

CAAGCAGAAGACGGCATACGAGAT<u>CTATCGCT</u>GTGACTGGAGTTCAGACGTGTGCTCTT

Reactions were transferred to a thermocycler programmed with the following stages: 95°C for 2min, 98°C for 30sec, 15 cycles of [98°C for 15sec, 64°C for 30sec, 72°C for 2min] (for pooled single nucleus reactions), 72°C for 5min, and 4°C forever. For bulk samples, we perform qPCR and subtract two cycles from the inflection point for each sample to determine optimal cycle number for amplification. PCR products were cleaned with two rounds of 0.8x SPRI beads, concentration was determined via Qbit® dsDNA High Sensitivity Assay Kit (Invitrogen Q32851), and library size was determined via Bioanalyzer (Agilent High Sensitivity DNA Kit, 5067-4626). Reactions were sequenced on an Illumina NextSeq 500 using the 300-cycle High Output v2 Kit or NovaSeq 6000 using the SP 300-cycle v1 kit.

### sNucDrop-seq library preparation and sequencing

FANS-sorted NeuN+ and NeuN- nuclei were individually diluted to a concentration of 100 nuclei/mL in DPBS containing 0.01% BSA. Approximately 1.25 mL of this single-nucleus suspension was loaded for each sNucDrop-seq run. The single-nucleus suspension was then co-encapsulated with barcoded beads (ChemGenes) using an Aquapel-coated PDMS microfluidic device (mFluidix) connected to syringe pumps (KD Scientific) via polyethylene tubing with an inner diameter of 0.38mm (Scientific Commodities). Barcoded beads were resuspended in lysis buffer (200 mM Tris-HCl pH8.0, 20 mM EDTA, 6% Ficoll PM-400 (GE Healthcare/Fisher Scientific), 0.2% Sarkosyl (Sigma-Aldrich), and 50 mM DTT (Fermentas; freshly made on the day of run) at a concentration of 120 beads/mL. The flow rates for nuclei and beads were set to 4,000 mL/hr, while QX200 droplet generation oil (Bio-rad) was run at 15,000 mL/hr. A typical run lasts 20 min. Droplet breakage with Perfluoro-1-octanol (Sigma-Aldrich), reverse transcription and exonuclease I treatment were performed, as previously described, with minor modifications^7^. For up to 120,000 beads, 200 μL of reverse transcription (RT) mix (1x Maxima RT buffer (ThermoFisher), 4% Ficoll PM-400, 1 mM dNTPs (Clontech), 1 U/mL RNase inhibitor, 2.5 mM Template Switch Oligo (TSO: AAGCAGTGGTATCAACGCAGAGTGAATrGrGrG) (Macosko et al., 2015), and 10 U/ mL Maxima H Minus Reverse Transcriptase (ThermoFisher)) were added. The RT reaction was incubated at room temperature for 30min, followed by incubation at 42C for 120 min. To determine an optimal number of PCR cycles for amplification of cDNA, an aliquot of 6,000 beads was amplified by PCR in a volume of 50 μL (25 μL of 2x KAPA HiFi hotstart readymix (KAPA biosystems), 0.4 μL of 100 mM TSO-PCR primer (AAGCAGTGGTATCAACGCAGAGT (Macosko et al., 2015), 24.6 μL of nuclease-free water) with the following thermal cycling parameter (95C for 3 min; 4 cycles of 98C for 20 sec, 65C for 45 sec, 72C for 3 min; 9 cycles of 98C for 20 sec, 67C for 45 sec, 72C for 3 min; 72C for 5 min, hold at 4C). After two rounds of purification with 0.6x SPRISelect beads (Beckman Coulter), amplified cDNA was eluted with 10 μL of water. 10% of amplified cDNA was used to perform real-time PCR analysis (1 μL of purified cDNA, 0.2 μL of 25 mM TSO-PCR primer, 5 μL of 2x KAPA FAST qPCR readymix, and 3.8 μL of water) to determine the additional number of PCR cycles needed for optimal cDNA amplification (Applied Biosystems QuantStudio 7 Flex). We then prepared PCR reactions per total number of barcoded beads collected for each sNucDrop-seq run, using 6,000 beads per 50- μL PCR reaction, and ran the aforementioned program to amplify the cDNA for 4 + 10 to 12 cycles. We then tagmented cDNA using the Nextera XT DNA sample preparation kit (Illumina, FC-131-1096), starting with 550 pg of cDNA pooled in equal amounts, from all PCR reactions for a given run. Following cDNA tagmentation, we further amplified the tagmented cDNA libraries with 12 enrichment PCR cycles using the Illumina Nextera XT i7 primers along with the P5-TSO hybrid primer (AATGATACGGCGACCACCGAGATCTACACGCCTGTCCGCGGAAGCAGTGGTATCAACGCAGAGT*A *C) (Macosko et al., 2015). After quality control analysis by Qubit 3.0 (Invitrogen) and a Bioanalyzer (Agilent), libraries were sequenced on an Illumina NextSeq 500 instrument using the 75-cycle High Output v2 Kit (Illumina). We loaded the library at 2.0 pM and provided Custom Read1 Primer (GCCTGTCCGCGGAAGCAGTGGTATCAACGCAGAGTAC) at 0.3 mM in position 7 of the reagent cartridge. The sequencing configuration was 20 bp (Read1), 8 bp (Index1), and 60 bp (Read2).

### Data Analysis

#### Read mapping and quality filtering for snhmC-seq and whole-genome bisulfite ACE-seq

The pre- processing (read alignment, quality filtering and read deduplication) was performed for snhmC-seq dataset of individual single cell as previously described with minor modifications^4^. Briefly, demultiplexing of inline barcodes was first performed allowing up to 1-nt mismatch. The data quality was examined with FastQC (http://www.bioinformatics.babraham.ac.uk/projects/fastqc/). Raw sequencing reads were trimmed for adaptor sequences and inline barcodes using Cutadapt^38^ with the following parameters in paired-end mode: -f fastq -q 20 -u 16 -U 16 -m 30 -a AGATCGGAAGAGCACACGTCTGAAC -A AGATCGGAAGAGCGTCGTGTAGGGA. The trimmed R1 and R2 reads were mapped independently against the reference genome (mm10) using Bismark^39^ (v0.18.2) with following parameters: --bowtie2 -D 15 -R 2 -L 20 -N 0 –score_min L,0,-0.2 (--pbat option was turned on for mapping R1 reads). Uniquely mapped reads were filtered for minimal mapping quality (MAPQ>=10) using samtools^40^. PCR duplicates were removed using the Picard *MarkDuplicates* (http://broadinstitute.github.io/picard/). To eliminate reads from strands not deaminated by A3A, reads with three or more consecutive non-converted cytosines in the CH context were removed using *filter_non_conversion* in Bismark. Low quality snhmC-seq datasets were removed using the following set of criteria. First, Mapping rate of raw sequencing reads is set to be at least 20%; Second, we set a minimum on the number of pass filter reads (uniquely mapped, deduplicated, MAPQ>=10) to be 100,000 per cell; Third, the 5mCG deamination rate in spike-in lambda phage control is higher than 90%. Overall, 360 out of 768 input nuclei passed our stringent quality filtering (47%). Base calling of unmethylated and methylated cytosines was performed by *bismark_methylation_extractor* in Bismark in each individual nucleus. 5hmC signals were calculated as % of C/(C+T) at each cytosine base. Sequencing reads for bisulfite ACE-seq were pre-processed as previously reported^17^.

#### Clustering analysis of single cell snhmC-seq

For **Fig. 1e**, 5hmCG methylation data were grouped into non-overlapping 100-kb bins across the whole genome of each nucleus. For each 100-kb bin, the 5hmCG level was computed by dividing the sum of methylated base calls by the sum of covered base calls. Due to the sparsity of the single-cell data, we adapted a previously reported imputation strategy^4^ for our snhmC-seq datasets. We imputed methylation rate at bins with coverage in 70% or more of the nucleus, replacing missing value with the average methylation across all cells for that bin. This allowed us to retain 76.9% (21,027/27,348) of 100-kb bins in the mouse genome for snhmC-seq data set. After imputation, Seurat 3 (v3.2.3)^41^ was used to perform the downstream clustering analysis. For normalization across different nuclei, methylation rate for all nuclei were scaled by library size (total methylation rate), multiplied by 10,000 and transformed to log scale using the function of *NormalizeData* in Seurat with the default parameter. The top 3,000 highly variable bins were identified and were then subjected into PCA analysis. The top 50 PCs were selected to construct a KNN graph using the function of *FindNeighors* in Seurat. Clusters were identified using the function FindCluster in Seurat. After examining the robustness of cluster classification with respect to several resolution and experimental parameters (cellular identity by FANS), we conservatively chose the resolution parameter of 1 to identify 3 cell clusters. For visualization, we performed dimensionality reduction using uniform manifold approximation and projection (UMAP), projecting all nuclei to two-dimensional space using the function *runUMAP* in Seurat.

To further explore the cell-type-specific gene’s 5hmCG level across cell types, we grouped 5hmCG methylation data into annotated genic region (from TSS to TES) and computed gene body methylation rate. We imputed the methylation rate of a gene with coverage in 60% or more of the nucleus, replacing low-coverage value (<5 CpG sites) with the average methylation across all cells for that gene. By doing this, 8,123 genes were kept, and the mean of their gene body 5hmCG methylation rate was calculated by defined cell types. (**Fig. 2b-c**).

#### Integration of snhmC-seq and snmC-seq datasets

To evaluate our snhmC-seq data in a neuronal subtype specific manner, we performed an integrated analysis of our data with snmC-seq datasets from mouse frontal cortical neurons^4^. Because 5hmCH is positively correlated with 5mCH across the neuronal genome (**Fig. S4e**, middle panel), we reasoned that the methylation signals of 5hmCH and 5mCH would allow joint analysis. For snmC-seq dataset, the base calling files (containing CG and CH methylation sites) were downloaded from GSE97179. CH methylation (5mCH+5hmCH) data were grouped into annotated genic regions. Because CH sites are more abundant than CG sites, we increased the coverage cutoff to >=100 CH site calls for this data set. We imputed the methylation rate of a gene with coverage in 90% or more of the nucleus, replacing low-coverage value (<100 CH sites calls) with the average methylation across all nuclei for that gene. 9,948 gene body 5mCH methylation levels of 3,377 nuclei were computed and imputed. For the snhmC-seq dataset, 262 nuclei assigned to two major neuronal clusters (Ex and Inh in **Fig. 1e**, right panel) were selected for integration analysis of neuronal methylome. For snhmC-seq, we lowered the coverage cutoff to >=25 CH sites. We imputed the methylation rate of a gene with coverage in 80% or more of the nucleus, replacing low-coverage value (<25 CH sites calls) with the average methylation across all nuclei for that gene. In the end, 10,782 genes of 262 neuronal nuclei were computed for the gene body’s 5hmCH methylation rate. For integration (**Fig. 2a**), we intersected the 2 gene lists from 5hmCH and 5mCH, and then kept the 9,685 common genes (97.4% of 5hmCH dataset; 89.8% of 5mCH dataset) for downstream analysis. Seurat 3 (v3.2.3) was used to perform joint analysis of 5hmCH and 5mCH. After internal normalization and identification of top 2,000 highly variable genes in Seurat, we identify anchors between 2 datasets using the 5mCH dataset as reference by the function of *FindIntegrationAnchors* with the dimensionality parameter of 20 in Seurat. The identified anchors were used for integration by the function of *IntegrateData* in Seurat. After integration, the integrated methylation level of the nuclei was scaled and centered for each gene across nuclei, and PCA analysis was performed on the scaled data. The 20 most significant PCs were selected and used for two-dimension reduction by UMAP. For visualization, the cellular identity from snmC-seq annotation or from snhmC-seq experiment was projected to the same UMAP.

#### Clustering and integration of single-nucleus RNA-seq (snRNA-seq)

FANS isolated NeuN+ and NeuN- nuclei were subjected to sNucDrop-seq for characterizing single-nucleus transcriptomes. The computational data analysis of sNucDrop-seq was performed as previously described with minor modifications ^27^. Briefly, after obtaining the digital expression matrices, *emptyDrops* function in R package DropletUtils (v1.2.2)^42^ was used to filter out the empty droplets. We used Scrublet (v0.2.1) ^43^ with the *expected_doublet_rate* parameter of 0.05 to identify the doublets which were removed. After QC steps with the parameters (nFeature_RNA > 500 & nFeature_RNA <= 3000 & percent.mt < 5) in Seurat (v3.2.3), 17,431 features across 6,918 nuclei were retained (**Fig. S6b**). Top 2,000 highly variable genes were identified. The expression level of highly variable genes was scaled and centered for each gene and was subjected to PCA. The top 30 PCs were selected and used for UMAP reduction in Seurat with the default parameters. Clusters were identified using the function *FindClustes* in Seurat with the resolution parameter set to 1. The marker genes were identified using FindAllMarkers in Seurat with the default parameters. The top-ranking markers of each cluster were used to annotate the clusters.

Since gene body CH methylation is inversely correlated with gene expression in mouse neuronal cells^4^. We reasoned that depletion of genic CH methylation would allow us to perform joint analysis between snmC-seq and sNucDrop-seq. For snmC-seq datasets, we computed the gene activity score (GAS):

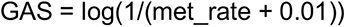

where met_rate represents the gene body CH methylation level.

After computing GAS for every gene across all single nuclei, we added GAS as a new assay in the Seurat object. After internal normalization and scaling steps, the function of *FindTransferAnchors* was used to identify the anchors between GAS and snRNA-seq by setting snRNA-seq dataset as a reference with the *reduction* parameter of “cca”. We imputed RNA expression into the snmC-seq nuclei based on computed anchors, then merged these 2 datasets into a combined Seurat object to perform co-embedding analysis in Seurat. PCA analysis was performed on the combined Seurat object. The first 30 PCs were selected and used for two-dimension reduction by UMAP (**Fig. S10b**). Clusters were identified using the function *FindClusters* in Seurat with the resolution parameter set to 1.

#### Genome browser visualization

We used Integrative Genomics Viewer (IGV, v2.3.91) to visualize bisulfite ACE-seq signals using mm10 Refseq transcript annotation as reference (upper panel in **Fig. 1d** and **Fig. S4d**). For **Fig. 1d**, 5hmCG signals (both strands combined) are indicated by upward ticks, with the height of each tick representing the fraction of modification at the site ranging from 0-50%.

#### Statistical analysis

Statistical analyses were performed using R. Statistical details for each experiment are also provided in the figure legends. No statistical methods were used to predetermine sample size for any experiments. All group results are expressed as mean +/− standard deviation unless otherwise stated. Specific p-values used for calling modified cytosine bases are explicitly stated in the text and figure legends. Each figure legend explicitly states the number of independent experiments.

#### Data visualization

All plots were generated using the ggplot2 (v3.3.0), cowplot (v1.0.0) and pheatmap (v1.0.12) packages in R (v3.5.1). In the box plots, the boxes display the median (center line) and interquartile range (from the 25th to 75th percentile), the whiskers represent 1.5 times the interquartile range and the circles represent outliers. In the violin plots, the gray line on each side is a kernel density estimation to show the distribution shape of the data; wider sections of the plot represent a higher probability, while the thinner sections represent a lower probability.

#### Data availability

All sequencing data associated with this study will be available on the NCBI Gene Expression Omnibus (GEO) database upon publication.

#### Code availability

The analysis source code underlying the final version of the paper will be available on GitHub repository (https://github.com/wulabupenn/snhmC-seq) upon publication.

